# Expanding the CRISPR toolbox by engineering Cas12a orthologs of metagenomic discovery

**DOI:** 10.1101/2024.09.27.615316

**Authors:** Dagang Tao, Bingrong Xu, Sheng Li, Hailong Liu, Suyu Shi, Yuan Wang, Changzhi Zhao, Jinxue Ruan, Liangliang Fu, Xingxu Huang, Xinyun Li, Shuhong Zhao, Shengsong Xie

## Abstract

Cas12a (Cpf1) is a CRISPR-associated nuclease with broad utility in genome editing and molecular diagnostic applications. However, the widespread adoption of CRISPR-Cas12a nucleases and their variants has been hindered by the requirement for a specific protospacer adjacent motif (PAM), relatively low CRISPR RNA (crRNA) activity and the inability to multiplex nucleic acid detection alone. To overcome these limitations, we employed a comprehensive framework combined with AlphaFold2 to *de novo* mine 1,261 previously unexploited Cas12a orthologs from the global microbiome. Following experimental analysis, we identified the most promising 21 Cas12a nuclease orthologs and designated them “Genie scissor 12” (Gs12). Our analysis uncovered two exceptional variants among these newly identified orthologs: Gs12-10, a first natural PAM-less Cas12a ortholog, which can recognize 52 distinct PAM types, representing a significant 1.8-fold expansion in recognition range compared to the relative LbCas12a PAM; and Gs12-7MAX, an engineered variant of Gs12-7 that exhibited 1.27-fold higher editing efficiency than enAsCas12a-HF. Furthermore, we harnessed Gs12-1, Gs12-4, Gs12-9, and Gs12-18, along with their corresponding engineered crRNAs, to develop a powerful four-channel multiplexed CRISPR-based nucleic acid detection system. The discovery of diverse functions in Cas12a offers a deeper understanding of the CRISPR/Cas12a family. Also, it holds great promise for expanding its applications and uncovering the untapped potential of other CRISPR/Cas systems.

**Graphical Abstract:** 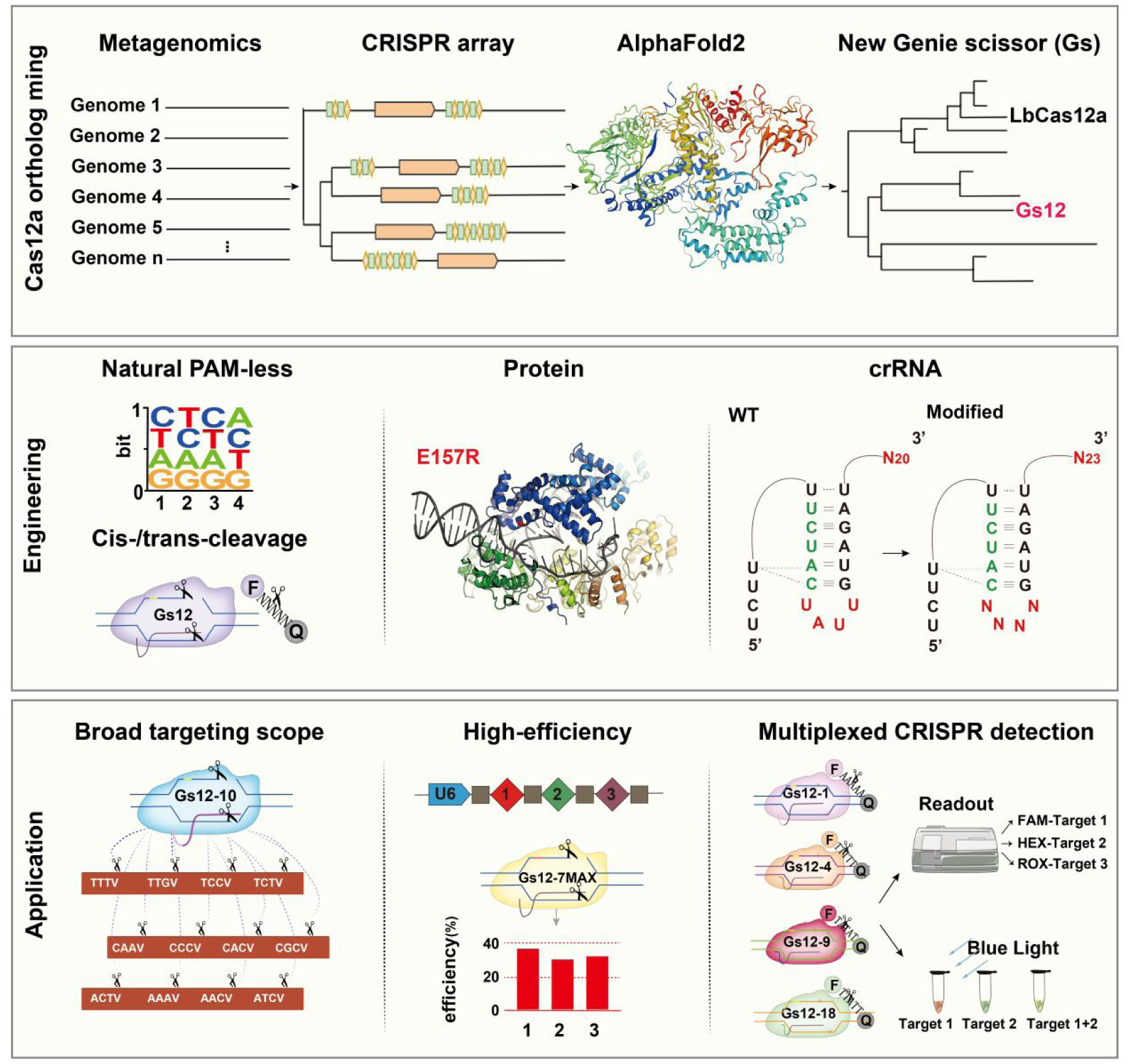

## Introduction

A diverse array of CRISPR effectors, including Cas12a (Cpf1), Cas9, Cas13 and Cas3, are widely employed as gene editing tools^1–6^. Notably, the Cas12 effectors, based on Class II type V proteins, exhibit distinct characteristics that distinguish them from Class II type VI and type II effectors^4^. These unique features include their ability to generate non-allelic DNA double-strand breaks (DSBs), compact structural design, and a single RuvC-like nuclease domain^7^. Leveraging these advantages, Cas12 family nucleases have garnered increasing attention and utilization in recent years^8^. Nevertheless, the distinct characteristics and applications of various subfamilies within the type V protein family remain understudied and lack a systematic understanding.

The CRISPR/Cas12a system is widely utilized for its ability to recognize the T-rich protospacer adjacent motif (PAM) at the 5’ end of its target and induce complementary interleaved DSBs within this spacer region^9^. Improved variants of Cas12a with enhanced activity and specificity have been developed. For example, an engineered Mb2Cas12a-RVRR variant has enabled editing with more lenient PAM requirements^10^. Engineered variants, such as enAsCas12a-HF1, have increased gene editing efficiency and reduced off-target effects^11^. Additionally, engineered CRISPR RNAs (crRNAs) with variable stem-loop designs can improve the adaptability of Cas12a^12,13^. Enhanced mammalian genome editing has also been achieved using new Cas12a orthologs with optimized crRNA scaffolds^14^.

Two primary bioinformatics-based methods are currently employed to discover CRISPR/Cas systems. One approach involves constructing hidden Markov models (HMMs) based on known Cas sequences, which are then applied to the analysis of bacterial and archaeal genomes, as well as metagenomic data derived from natural environments^15,16^. The other method involves comparing the hallmark sequences characteristic or family-specific features of CRISPR/Cas systems with the genomes of bacteria and archaea using PILER-CR or a machine learning approach^17,18^. This has enabled the discovery of new Cas12a orthologs through *de novo* mining of prokaryotic genomes^19^. Several examples of discovered Cas12a orthologs include a study of genome editing activity for 16 Cas12a orthologs^20^, characterization of Cme and Yme thermostable Cas12a orthologs^21^, and the discovery and engineering of AiEvo2, a novel Cas12a nuclease for human gene editing applications^22^. However, obtaining complete and high-quality genomic data for bacteria and archaea and developing well-established data mining methodologies remains a significant challenge, hindering the discovery of new Cas12a variants.

Notably, by engineering the CRISPR-optimized AsCas12a (opAsCas12a) system with critical modifications to the Cas nuclease and its crRNA, high-efficiency combinatorial genetic screening can be achieved, as demonstrated by Gier *et al.*^23^. Researchers have shown the use of Cas12a for simultaneous editing of multiple genes and sites^10,24^. Moreover, novel detection platforms have been developed by exploiting the collateral activities of Cas12 nuclease, which selectively cleaves single-stranded DNA (ssDNA)^12,25^. Previous studies reported a CRISPR-Cas12-based detection method for SARS-CoV-2^26^. Furthermore, a one-pot isothermal Cas12-based assay for sensitively detecting microRNAs was developed^27^. A naked-eye colorimetric detection method for nucleic acids based on CRISPR/Cas12a and a convolutional neural network was also developed^28^. Moreover, a naked-eye CRISPR-Cas12a/Cas13a multiplex point-of-care detection of genetically modified swine was developed^29^. Simultaneous breast cancer biomarkers circROBO1 and BRCA1 detection based on a CRISPR-Cas13a/Cas12a system have been achieved^30^. However, developing CRISPR-based multiplex nucleic acid detection platforms by using the same Cas orthologs remains a significant technological challenge due to the scarcity of suitable Cas12a nucleases.

In recent years, the applications of Cas12 effectors for gene editing in animals and plants have rapidly expanded, including the development of base editors, primer editor tools for regulating gene expression, methods for gene targeting, and biosensors^12,31,32^. Moreover, these exciting advancements have sparked new ideas for manipulating, modifying, and detecting DNA molecules, potentially paving the way for precise and efficient CRISPR-Cas12a-based gene editing in the future^33,34^. However, the development of CRISPR-Cas12a remains constrained by several limitations, including its low enzymatic activity, PAM restrictions, and the lack of specific ssDNA cleavage preferences^5,35^. Therefore, expanding our repertoire by discovering additional Cas12a family members from natural bacterial sources is essential to enriching the CRISPR-Cas12a toolbox and overcoming these limitations.

This study sought to identify novel Cas12a systems exhibiting high *cis*- and *trans*-cleavage activities, broad PAM range, or specific ssDNA preference. To achieve this, we explored the evolutionary diversity of CRISPR/Cas12a systems through a combination of bioinformatic screening and characterization. Our screen of 1261 Cas12a systems identified 21 active enzymes *in vitro*, including two that demonstrated activity in mammalian cells. We introduced the Genie scissor 12 (Gs12) series nomenclature for our newly discovered enzymes to facilitate easy identification and distinction from other RNA-guided endonucleases. The discovery of diverse functions in Gs12 endonucleases provides a deeper understanding of the CRISPR/Cas12a family and the potential to unlock new applications and uncover the hidden capabilities of other CRISPR/Cas systems.

## Results

### Identification and comprehensive evaluation of the new Cas12a orthologs for genetic engineering

To expand the targeting scope and enhance the *cis* or *trans-*cleavage activity of Cas12a, we developed a computational pipeline to identify novel naturally occurring Cas12a orthologs (Figure 1A). We started by filtering 278,629 genome bins larger than 20 kb from the Global Microbial Gene Catalog (GMGC)^36^ to detect genomes containing CRISPR-Cas structures. Next, we used the MinCED^37^ program to screen the CRISPR arrays and retrieved open reading frames using Prodigal ^38^. In our search for novel Cas12a nucleases suitable for genome editing, we employed PSI-BLAST^39^ and identified 1,261 CRISPR-Cas12a loci not previously utilized for genome editing (Figure S1A). Phylogenetic tree analysis revealed that the newly discovered candidates belonged to distinct Cas12a subtypes, exhibiting evolutionary branches separate from those of Cas12b, Cas12c, Cas12f, and Cas9, suggesting their potential classification as new members of the Cas12a subfamily (Figure 1B and Figure S1B). Additionally, our analysis revealed that the majority of Type V Cas12a locus systems consist of Cas1, Cas2, Cas4 and a CRISPR array (Figure S2).

**Figure 1.**
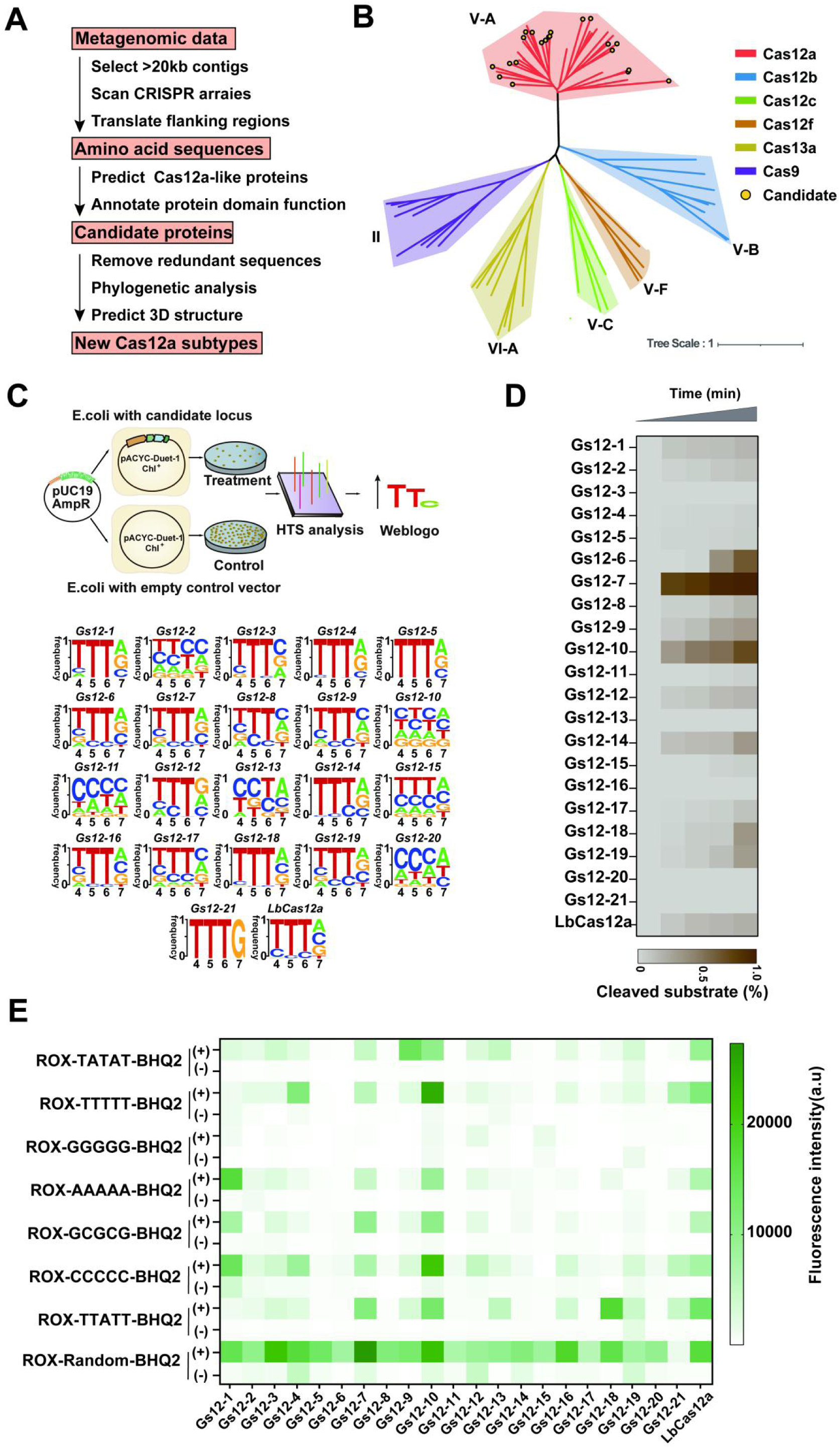
Identification of novel CRISPR-Cas12a systems. **A.** Flowchart depicting the bioinformatics pipeline used in identifying novel Cas12a families. **B.** Phylogenetic tree of 73 Cas protein sequences. The branches of the tree corresponding to the 21 candidates’ Cas12a orthologs examined in this study are marked with a dark circle. **C.** Schematic workflow representation to determine PAM sequence requirements for dsDNA cleavage. Twenty-one Cas12a candidates are referred to as Gs12-1 to Gs12-21 herein. Time from 30 s, 2 min, 10 min, 20 min. HTS, High Throughput Sequencing. **D.** Cleavage activity of Cas12a orthologs was measured using *in vitro* DNA cleavage assays using dsDNA substrates. **E.** Evaluation of *trans* nuclease substrate preferences for Cas12a orthologs using ssDNA-FQ reporters. ROX, which stands for carboxy-X-rhodamine, is a fluorescent compound with an excitation peak at 576 nm and an emission peak at 601 nm. BHQ2, Black hole quencher-2.

To enhance the predictive accuracy of Cas12a orthologs, we employed Alphafold2^40^ to predict the three-dimensional structures of 21 candidate nucleases (Figure S3). We performed structural comparisons between known and novel Cas12a nuclease pairs using TMalign^41^, and established a threshold TM-score greater than 0.7 to identify structurally similar Cas12a orthologs. These results indicated that the spatial structure of candidate Cas12a orthologs is closely related to known Cas12a proteins (Figure S4 and S5A). Subsequently, we searched the Cas12a candidates for novel ortholog sequences with less than 60% identity, aiming to narrow down further the 21 candidates (herein referred to as Gs12-1 to Gs12-21) (Figure S5B). Next, a comparative analysis of the two key structural domains, RuvC and PI, revealed significant differences in their amino acid compositions among known Cas12a proteins (Figure S6 and S7). Moreover, the alignment of the Gs12-1 to Gs12-21 mature crRNA sequences revealed the utilization of 7 highly conserved crRNA scaffolds and their corresponding loop regions (Figure S8). The following different loop sequences were found: “AUU”, “GU”, “GUU”, “CUC”, “UAGU”, “UUC”, and “CUU”. However, sequence variations were observed most in the loop region and only one in the stem region (Figure S8).

We assessed the PAM profiles and *cis*- and *trans*-cleavage activities of the 21 orthologs from our collection. These Gs12 genes encode approximately 1200 to 1300 amino acids, with the sequences corresponding to crRNA synthesized (Supplementary Table 1). Subsequently, the Gs12 nucleases expressed in *Escherichia coli* (*E. coli*) cells were purified and subjected to DNA cleavage assays, in which they were incubated with *in vitro*-transcribed crRNAs and dsDNA substrates. To characterize the global PAM specificity profiles of these Gs12 orthologs, we employed a bacterial-based negative selection system analogous to the methods previously utilized for identifying PAM preferences of Cas9 (Figure 1C)^42^. This entailed a bacterial PAM-depletion assay, in which a library of plasmids bearing seven randomized base pairs adjacent to a protospacer was subjected to cleavage by Gs12 in *E. coli*. The plasmids containing the preferred PAMs were depleted, and the remaining plasmids were sequenced and analyzed.

We generated profiles of PAM specificity for these Gs12 candidates (Figure 1C, Supplementary Table 2). It is noteworthy that Gs12-1 exhibited a substantial depletion of sites with a 5’-“HTTV” (H: A/C/T; V: A/C/G) PAM, whereas Gs12-3 showed a higher specificity towards a 5’-“BTTV” (B: G/T/C; V: A/C/G) PAM. Conversely, Gs12-2 demonstrated the capacity to recognize a simple 5’-“NNHN” (H: A/C/T; N: A/T/G/C) PAM for dsDNA cleavage (Figure 1C). Except for Gs12-2, Gs12-10, Gs12-11, Gs12-13, and Gs12-20, the predominant PAM preference among other Gs12 variants was the canonical 5’-“TTTV” (V: A/C/G) like that of LbCas12a (Figure 1C). Notably, Gs12-10 exhibited no specific PAM sequence restriction, suggesting it might function as a first natural PAM-less Cas12a ortholog (Figure 1C).

Furthermore, we investigated the dsDNA cleavage activities of the 21 purified Gs12 nucleases *in vitro* using dsDNA substrates containing 20 nucleotide (nt) target sequences with the canonical “TTTV” PAM. LbCas12a was included as a positive control. Our results showed that these purified Cas12a orthologs recognized the 5’-“TTTV” PAM to cleave target dsDNA, albeit with different efficiencies (Figure 1D). Notably, Gs12-7 exhibited complete cleavage of dsDNA substrates *in vitro*, whereas Gs12-6, Gs12-9, Gs12-12, Gs12-10, Gs12-14, and Gs12-19 exhibited partial cleavage of target dsDNA substrates, mirroring observations with LbCas12a (Figure 1D). In addition, we assessed the *trans*-cleavage activity of the purified Gs12 nucleases using ssDNA-FQ reporters with varying base compositions. Our results show that most purified Gs12 orthologs recognized the ssDNA-FQ reporter with random bases (ROX-random-BHQ2) configuration, similar to LbCas12a (Figure 1E). However, Gs12-4 and Gs12-10 strongly preferred polyT (ROX-TTTTT-BHQ2) (Figure 1E). Notably, Gs12-1 displayed a remarkable preference for polyA (ROX-AAAAA-BHQ2), while Gs12-9 showed high affinity for TA repeats (ROX-TATAT-BHQ2) and Gs12-18 showed high affinity for ROX-TTATT-BHQ2 (Figure 1E). These results highlight the different PAM profiles and distinct *cis-* or *trans*-cleavage activities observed among the newly identified Cas12a orthologs.

### PAM-less genome editing using a CRISPR–Gs12-10 toolbox

We then investigated the ability of Gs12-10 to generate DSBs at various locations harbouring different PAMs along a plasmid or linear DNA substrate (Figure 2A). To assess its *in vitro* activity, we performed digests against 16 distinct target sites on a circular plasmid substrate, spanning “ANTN” PAMs. The results revealed that Gs12-10 efficiently cleaved all these sites to near completion (Figure 2B). To further investigate the PAM-less nature of Gs12-10, we performed a comprehensive comparison between LbCas12a and an additional 64 crRNAs targeting different target sites encompassing all possible combinations of the 2nd, 3rd, and 4th positions within an “NNN” PAM (Figure 2C). Consistent with previous studies^4,20^, LbCas12a exhibited robust substrate cleavage activity when programmed with crRNA targeting sites with “TTTV” PAMs. In addition, sporadic activity against sites with “VTTV”, “TCTV”, or shifted “TCCV” PAM sequences was observed *in vitro* (Figure 2C). In contrast, Gs12-10 efficiently cleaves the same substrates as LbCas12a and achieves near-complete digestion of additional substrates. These include sites containing “AGAV”, “ATGV”, “AGCV”, etc., PAMs (Figure 2C).

**Figure 2.**
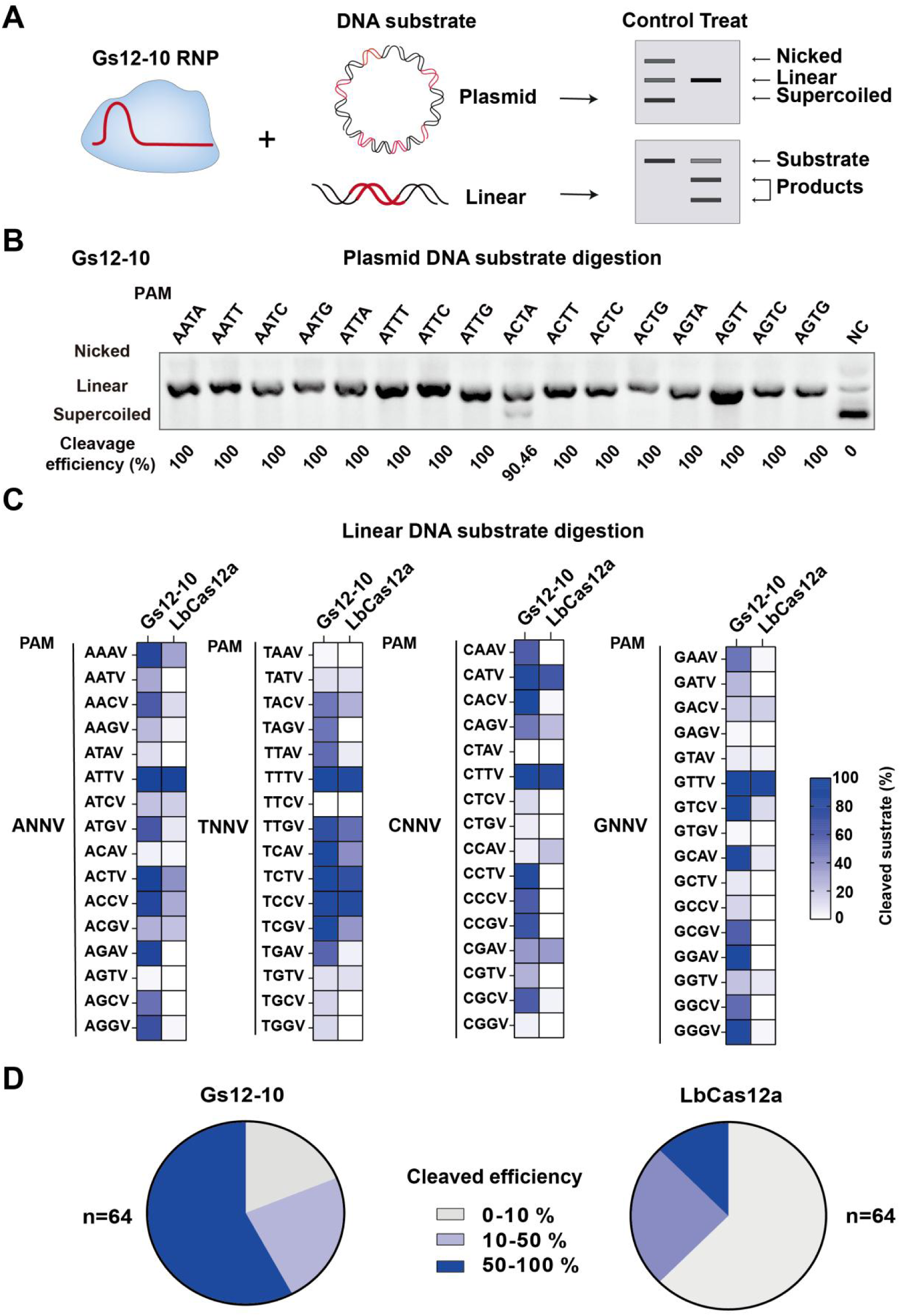
Characterization of Gs12-10 *in vitro* cleavage efficiencies. **A.** Schematic representation of the DNA *in vitro* cleavage efficiencies for Gs12-10. **B.** Evaluation of the cleavage efficiencies of Gs12-10 on circular plasmids targeting the “ANTN” PAM. **C.** Comparison of the *in vitro* cleavage efficiencies of Gs12-10 and LbCas12a across 64 target sites representing all 1st, 2nd or 3rd position combinations of an “NNNV” PAM. PAM (protospacer adjacent motif) refers to a specific sequence pattern, where N represents any of the four nucleotide bases A, T, C, or G, and V represents any of the three nucleotide bases A, C, or G. **D.** Summary of the proportion of crRNAs that led to near-complete (50-100%), partial (10-50%) or incomplete (0-10%) substrate cleavage when using Gs12-10 and LbCas12a. Mean is shown for n = 3.

Notably, Gs12-10 achieved nearly complete substrate digestion with 37 of the 64 crRNAs, partial digestion with 15 crRNAs, and low or no activity with the remaining 12 (Figure 2D). Conversely, LbCas12a showed nearly complete substrate digestion with only 8 of 64 crRNAs, partial digestion with 21 crRNAs, and low or no activity with the remaining 35 (Figure 2D). Overall, Gs12-10 effectively cleaved 81% (52/64) of the PAM target sequences tested, whereas LbCas12a showed cleavage activity with only 45% (29/64) of the PAM sequences. This result showed that Gs12-10 exhibits an unprecedented ability to recognize a diverse range of PAMs. Specifically, Gs12-10 recognizes 52 distinct PAM types, representing a 1.8-fold increase in the range of recognition compared to the relative LbCas12a PAM. This finding further suggests that Gs12-10, unlike other Cas12a orthologs, functions as a nearly PAM-independent nuclease with exceptionally high cleavage efficiency.

### Genome editing in mammalian cells with PAM-less Gs12-10

We constructed a eukaryotic expression plasmid containing the synthetic codon-optimised candidate to demonstrate the extended targeting range of Cas12a ortholog by Gs12-10. We evaluated its activity against endogenous genes in human cells (Figure S9A). First, we assessed the editing efficiency of Gs12-10 in HEK293T cells by targeting three genes - FANCF, RUNX1, and EMX1 - using 20 base pair (bp) spacers against sites containing the canonical “TTTV” PAM. The results revealed that Gs12-10 exhibited an editing efficiency comparable to LbCas12a’s (Figure S9B). We then sought to enhance the editing activity of Gs12-10 by optimizing the spacer length in the range of 20-25 bp. Our results showed that crRNAs with 22 and 23 bp spacer lengths exhibited significantly higher editing activity than those with 20 bp spacers (Figure 3A). Furthermore, by comparing the editing efficiencies targeting DNMT1 and FANCF, we observed that crRNAs with a length of 23 nt outperformed those with a length of 22 nt. Therefore, we selected the optimal spacer length of 23 bp for subsequent experiments (Figure 3A).

**Figure 3.**
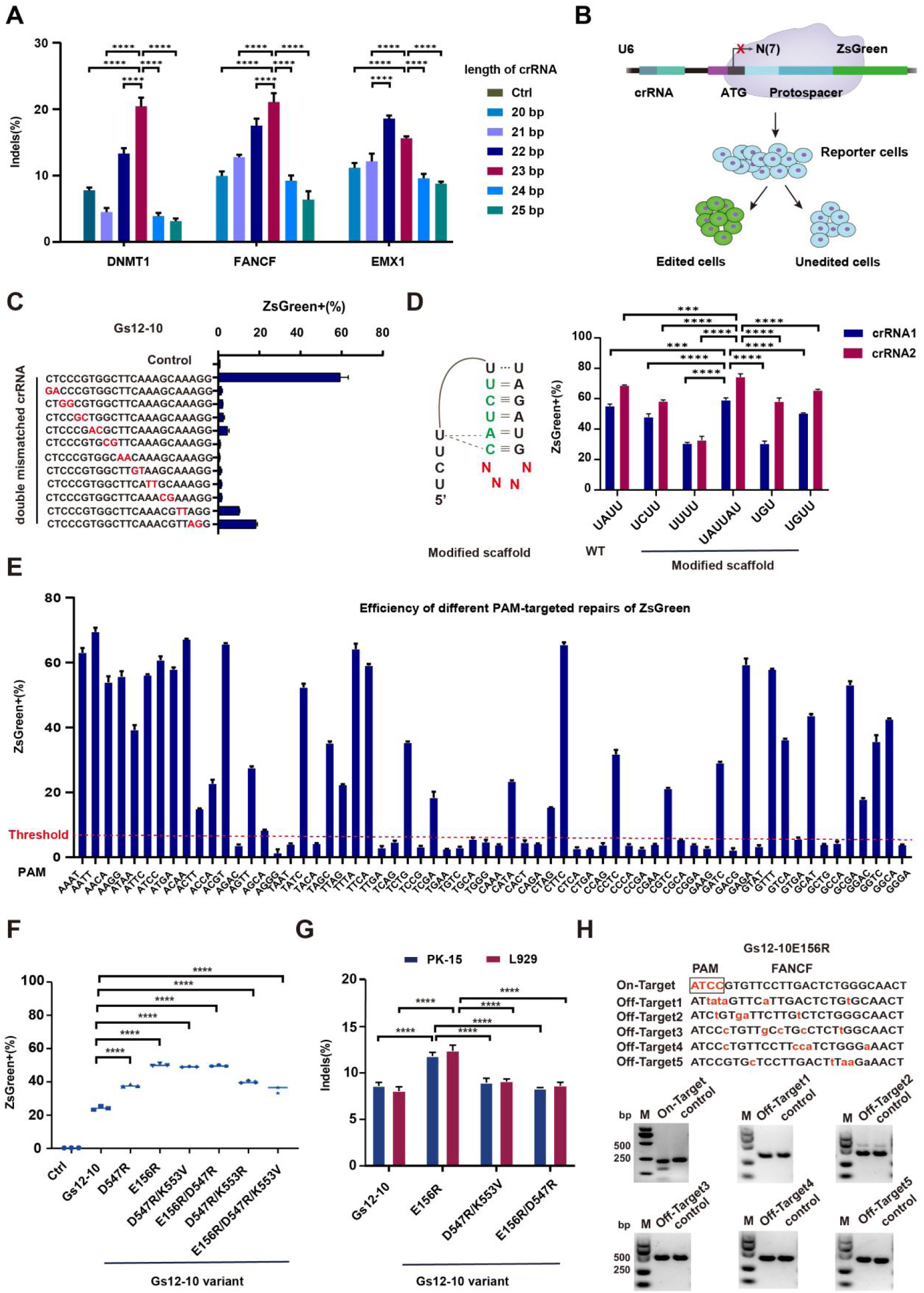
Enhance the editing activity of Gs12-10 and characterized its PAM-less features in mammalian cells. **A.** Enhanced targeted Indels frequencies by optimized spacer length. Indels stands for insertions and deletions, Ctrl stands for control group, bp stands for base pair, **** P< 0.0001. **B.** Schematic representation of the DNA cleavage efficiencies Gs12-10 as measured by the ZsGreen activation assay. N(7) represents seven random nucleotide bases, which can be any of the four bases A, T, C, or G. **C.** Evaluation of the cleavage ability of Gs12-10 for recognize two mismatched target sites using the ZsGreen activation assay. The red letters indicate the presence of two mismatched bases. **D.** Enhanced targeted Indels frequencies by optimized crRNA scaffold using the ZsGreen activation assay. The red letters “N” indicated the presence of modified bases in the loop sequence, WT stands for Wild-Type scaffold, \*\*\**P*< 0.001, **** *P*< 0.0001. **E.** Evaluation of the editing activity of Gs12-10 in human cells using the ZsGreen activation assay at 64 target sites that represent all possible combinations of “NNNV” PAM sequences at the 1st, 2nd, and 3rd positions. PAM stands for protospacer adjacent motif. Note: The term “Threshold” refers to the relatively lower level of editing activity (< 8%). **F.** Modification of the Gs12-10 nuclease through alternative amino acid substitutions enhances editing activity as measured by the ZsGreen activation assay. **** *P*< 0.0001. **G.** Gs12-10 variants enhance on-target editing activity in pig and mouse cells. PK-15 stands for the pig-derived PK-15 cell line. L929 stands for the mouse-derived fibroblast L929 cell line. **** *P*< 0.0001. **H.** Detection of predicted off-target endogenous sites of Gs12-10E156R (Gs12-10MAX) containing the “ATCC” PAM by using T7E1 cleavage assay. Mean is shown for n = 3.

We then used a green fluorescent protein (ZsGreen) activation assay to assess the activity of Gs12-10 in human cells, with modifications as previously described^43^. In this assay, Cas12a-mediated genome editing results in insertions or deletions (Indels), leading to ZsGreen expression in a subset of cells (Figure 3B). Targets with two base mismatches to the crRNA were generated to evaluate specificity. The results showed that the Gs12-10 nuclease decreased ZsGreen-positive cells, which can moderately tolerate the mismatches, similar to LbCas12a (Figure 3C, Figure S10). To further increase the editing efficiency of Gs12-10, we optimized the loop structure within the crRNA backbone. Through the ZsGreen activation assay, we observed that the addition of two bases, “AU”, to the wild-type “UAUU” loop with two different spacers improved the editing efficiency compared to the wild-type scaffold (Figure 3D). The DNA cleavage activity of 64 PAM profiles of Gs12-10 using optimized crRNAs in human HEK293T cells by ZsGreen activation assay was investigated. We found that 35 of 64 crRNAs have relatively high cleavage activity (Figure 3E). The tested sites containing non-canonical PAMs, such as “AACV”, “ACCV”, “AGGV”, etc., allowed for the repair editing of the ZsGreen protein by Gs12-10. Next, we also tested 20 randomly selected endogenous target sites. The results showed that all these sites, which contained non-canonical PAMs, could be edited (Figure S11).

To generate a highly active and specific Gs12-10 mutant, we constructed two variants with single amino acid substitutions (E156R, D547R) and additional variants with combinations of these substitutions (E156R/D547R, D547R/K553R, D547R/K553V, and E156R/D547R/K553V)^11^. All six variants were evaluated in human HEK293T cells and showed increased gene editing activity at sites with canonical “TTTV” PAMs compared to wild-type Gs12-10 as determined by the ZsGreen activation assay (Figure 3F). Among these variants, the E156R mutation exhibited the highest editing activity in human cells and in porcine PK-15 and murine L929 cells (Figure 3G). Based on these results, we designated this optimized variant (E156R) as Gs12-10MAX. The fidelity of the Gs12-10MAX variant was then evaluated against that of LbCas12a. Targets with two base mismatches to the crRNA were generated for comparison. The results showed that the Gs12-10MAX variant nuclease induced a decrease in ZsGreen-positive cells compared to its wild-type Gs12-10 nuclease and LbCas12a (Figure 3C, Figure S10 and Figure S12). We then predicted potential off-target sites and performed off-target analyses of the crRNAs on two endogenous targets (DNMT1 and FANCF) with canonical “TTTG” and non-canonical “ATCC” PAMs in human cells. The results showed no detectable off-target effects for crRNAs (Figure 3H and Figure S13). These results indicate that the engineered Gs12-10MAX variant exhibits high specificity in genome editing.

### Highly efficient genome editing in mammalian cells with Gs12-7 and variants

Among the newly identified Gs12 nucleases, Gs12-7 exhibited the highest *in vitro* cleavage activity (Figure 1D). To further explore the capabilities of this enzyme, we evaluated its gene editing activity in human cells using two distinct strategies: either delivering pre-assembled Ribonucleoprotein (RNP) complexes consisting of Gs12-7 nuclease and crRNA or co-expressing Gs12-7 and crRNA from plasmids (Figures 4A and 4B). The results showed that Gs12-7 achieved an average activity of nearly 20%. However, compared to enAsCas12a-HF1, the activity of Gs12-7 was relatively low (Figure 4C). To engineer a highly active and specific Gs12-7 mutant, we generated two variants: one with a single amino acid substitution (E157R) and another with three individual substitutions (E157R, G516R, K522V) (Figure 4D). We evaluated these variants in human HEK293T cells by targeting individual sites. We found that the Gs12-7E157R variant exhibited significantly enhanced gene editing activity at sites with canonical “TTTV” PAM compared to enAsCas12a-HF1 (with substitutions E174R/N282A/S542R/K548R of AsCas12a), as measured by the high-throughput sequencing (Figures 4E and 4F). Notably, when editing three or two target sites simultaneously, the Gs12-7E157R variant demonstrated even higher gene editing activity than enAsCas12a-HF1 and another Gs12-7 variant (Figures 4G and 4H). Our results indicate that the engineered variant Gs12-7E157R exhibits a significant 1.27-fold increase in editing efficiency compared to enAsCas12a-HF1. Notably, even without employing sorting methods for positive cells, the cleavage activity of Gs12-7E157R reaches an impressive level of up to 42% (Figure 4). Based on these findings, we have designated this optimized variant (E157R) as Gs12-7MAX, highlighting its potential as a powerful tool for precision genome editing applications. Based on these results, we designated this optimized variant (E157R) as Gs12-7MAX. Subsequently, we tested the outcome of gene editing for CRISPR-Gs12-7MAX.

**Figure 4.**
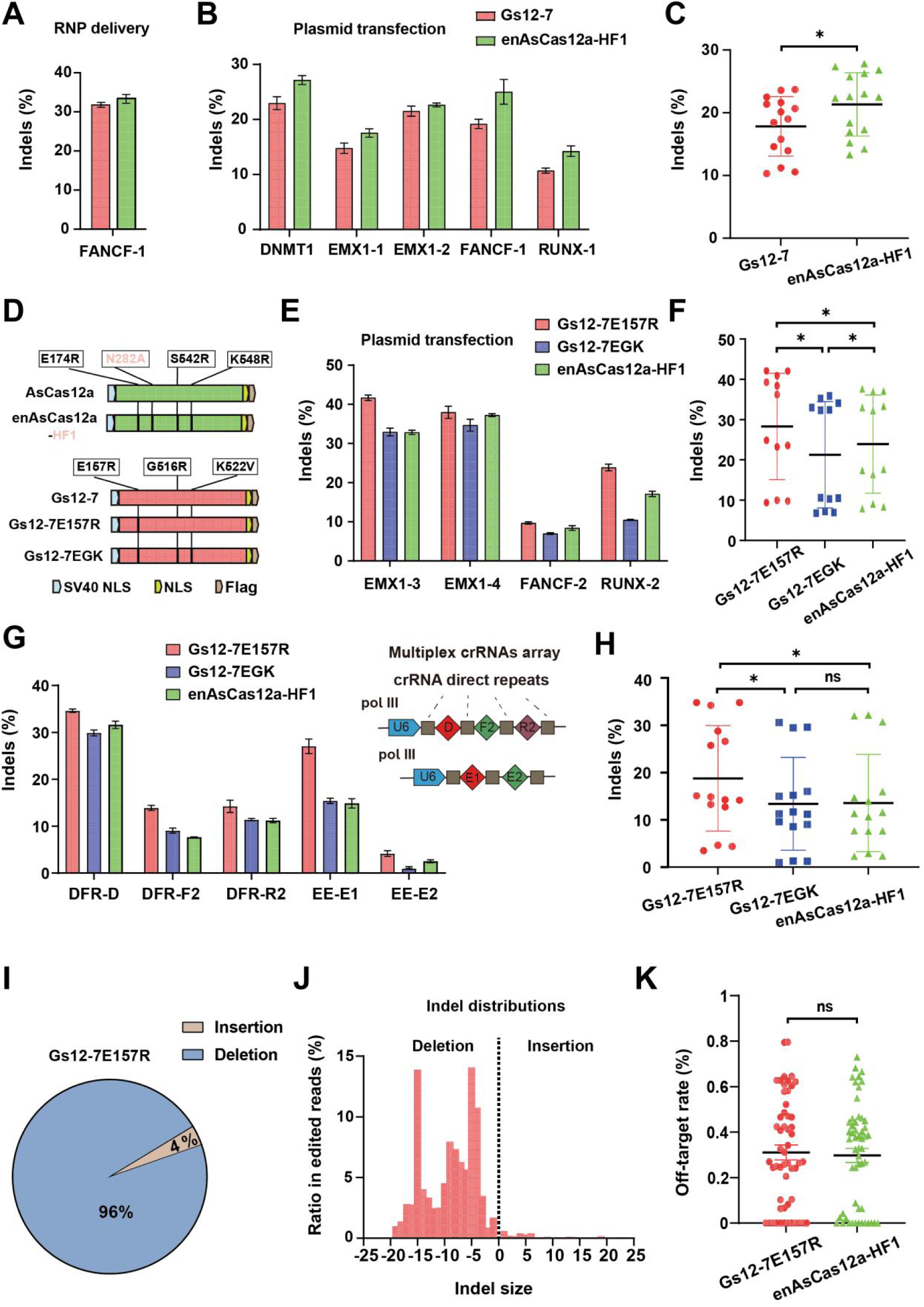
Evaluation of Gs12-7 and its variants’ activity and specificity. **A.** Detection of the activity of Gs12-7 using RNP delivery. RNP (ribonucleoprotein) refers to a complex consisting of the Gs12-7 protein and crRNA. **B-C**. Comparison of the activity of Gs12-7 and enAsCas12a-HF1 using the plasmid transfection method. Indels stands for insertions and deletions, * *P*<0.05. **D.** Pattern diagram of Gs12-7 variants and enAsCas12a-HF1. NLS, stands for nuclear localization signals, * *P*<0.05. **E-F.** Comparison of the activity of two Gs12-7 variants (Gs12-7E157R and Gs12-7EGK) and enAsCas12a-HF1 for targeting a single site. * *P*<0.05. **G-H**.Comparison of the activity of two Gs12-7 variants and enAsCas12a-HF1 for simultaneous targeting of two or three sites. DFR stands for a set of three gene targets: DNMT1, FANCF, and RUNX1. EE stands for two target sites of the EMX1 gene. D: DNMT1, F2: FANCF, R2: RUNX1, E1: EMX1-site1, E2: EMX1-site2, Pol III stands for U6 Pol III promoters, * *P*<0.05, ns, non-signficant. **I-J.** Detecting the outcome of Gs12-7E157R.The types of gene editing (I) and Indel distributions (J) of Gs12-7E157R. **K.** Comparison of the specificity of Gs12-7E157R and enAsCas12a-HF1. ns non-significant.

The results showed that the ratio of 96 % genotype was deletions, ranging in size from 0 to 20 bp, and 4 % genotype was insertions (Figure 4I and 4J), similar to Gs12-7EGK and enAsCas12a-HF1 (Figure S14A and S14B). We then predicted 18 potential off-target sites and performed off-target analyses of the crRNAs on six endogenous targets using high-throughput sequencing in human cells. The results showed that the off-target effects between Gs12-7MAX and enhanced high-fidelity enAsCas12a-HF1 were almost identical (Figure 4K and Figure S15). These results indicate that the engineered Gs12-7MAX variant exhibits high specificity in genome editing.

### CRISPR/Gs12s-mediated multiplexed nucleic acid detection

To address the diverse requirements for simultaneous detection of multiple targets, we developed a multiplex CRISPR detection platform that exploits the distinct *trans*-cleavage preferences of Gs12s enzymes (Figure 1E). In our quest to identify suitable candidate enzymes for multiplex detection, we performed further biochemical characterization on four members of the CRISPR-Gs12 family: Gs12-1, Gs12-4, Gs12-9, and Gs12-18 (Figure 5A). *Trans*-cleavage preferences were further evaluated using four ssDNA-FQ reporters (Figure 5A). Our results demonstrate that Gs12-1 exhibits a preference for the motif “ROX-AAAAA-BHQ2”, Gs12-4 for “ROX-TTTTT-BHQ2”, Gs12-9 for “ROX-TATAT-BHQ2”, and Gs12-18 for “ROX-TTATT-BHQ2” (Figure 5B).

**Figure 5.**
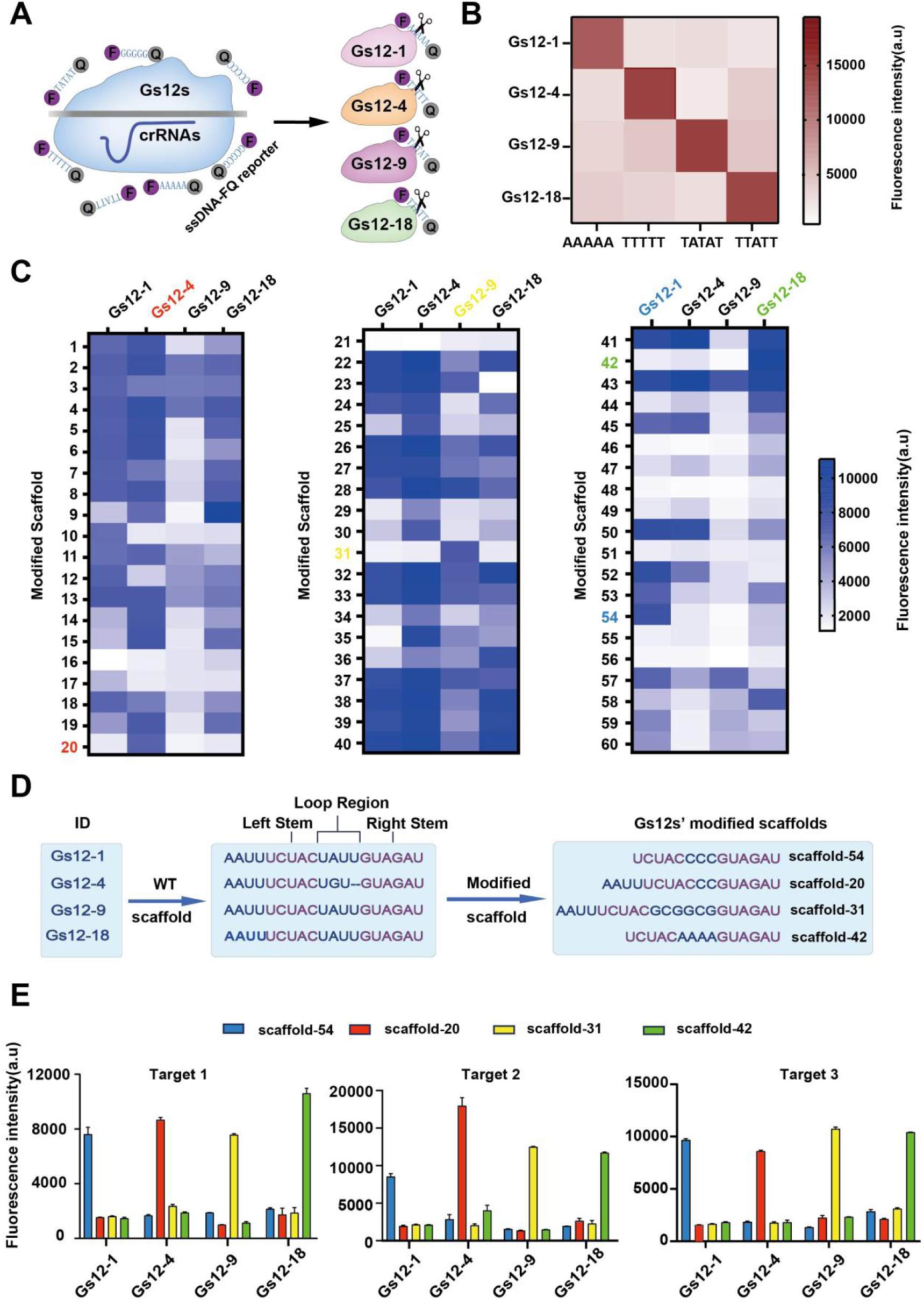
Identification of Gs12 nucleases with corresponding ssDNA-FQ reporter preferences and engineered crRNA for four-channel multiplex nucleic acid detection. **A-B.** Evaluation of the ssDNA-FQ reporters preference of Gs12-1, Gs12-4, Gs12-9, and Gs12-18 nucleases. ssDNA-FQ stands for single-stranded DNA Fluorescence-Quenching reporter, F: Fluorescence, Q: Quenching. **C.** Screening and identification of specific scaffold variants in crRNAs to improve the corresponding detection specificity of Gs12-1, Gs12-4, Gs12-9, and Gs12-18 nucleases. a.u stands for arbitrary unit. **D.** Identification of corresponding engineered crRNAs for Gs12-1, Gs12-4, Gs12-9, and Gs12-18 nucleases. ID stands for Identity, WT stands for Wild-Type crRNA. **E.** Evaluation of the detection ability of CRISPR-Gs12-based four-channel multiplex nucleic acid detection assay. Target 1 refers to the p72 gene of African Swine Fever Virus. Target 2 refers to the VP1 gene of Seneca Valley Virus. Target 3 refers to the E2 gene of Classical Swine Fever Virus. Mean values are shown for n = 3.

Subsequently, the *trans*-cleavage specificity of Gs12-1, Gs12-4, Gs12-9, and Gs12-18 was evaluated against the same target site p72 gene of African Swine Fever Virus (ASFV). In particular, their respective structurally engineered scaffolds of crRNAs were also assessed. A total of 60 different types of engineered scaffolds were compared, and the results showed that the fluorescence intensity varied depending on the combination of four kinds of Gs12s and crRNAs used (Figure 5C). Among them, we identified four modified scaffolds -54, 20, 31, and 42, corresponding to Gs12-1, Gs12-4, Gs12-9 and Gs12-18 enzymes, respectively, allowing for specific target detection (Figures 5C and D). Next, we established different comparison groups using three target sites: Target 1, the p72 gene of ASFV; Target 2, the VP1 gene of Seneca Valley Virus (SVA); and Target 3, the E2 gene of Classical Swine Fever Virus (CSFV). We also used the same ssDNA-FQ reporter but with crRNAs of structurally engineered scaffold variants as follows: Gs12-1, Gs12-4, Gs12-9 and Gs12-18 paired with crRNAs of scaffold-54, scaffold-20, scaffold-31 and scaffold-42, respectively, to detect these three target site. The results showed that Gs12-1, Gs12-4, Gs12-9 and Gs12-18, each using their respective structurally engineered crRNA of scaffold-54, scaffold-20, scaffold-31 and scaffold-42, were able to identify their respective target molecules (Figure 5E) specifically. These results show that we have established a novel four-channel multiplex nucleic acid detection platform using four different Cas12a orthologs and their corresponding structurally engineered crRNAs and ssDNA-FQ reporters.

### A multiplex Recombinase polymerase amplification (RPA) coupled with CRISPR-Gs12 system for highly sensitive and specific detection of nucleic acid

We evaluated the CRISPR/Gs12s-mediated nucleic acid detection system as a universal platform for multiplexed nucleic acid detection by demonstrating its application in detecting ASFV, SVA, and CSFV. The operating principle of this multiplexed detection platform involves the use of three Gs12 nuclease, each with corresponding structurally engineered scaffold of crRNA and ssDNA-FQ reporters, including Gs12-1/scaffold-54/FAM-AAAAA-BHQ1, Gs12-4/scaffold-20/HEX-TTTTT-BHQ1, and Gs12-18/scaffold-42/ROX-TTATT-BHQ2 (Figure 6A). We used plasmids containing ASFV p72, SVA VP1, and CSFV E2 as detection templates and found that the CRISPR/Gs12s-mediated nucleic acid detection system could clearly distinguish between single or mixed targets (Figure 6B).

**Figure 6.**
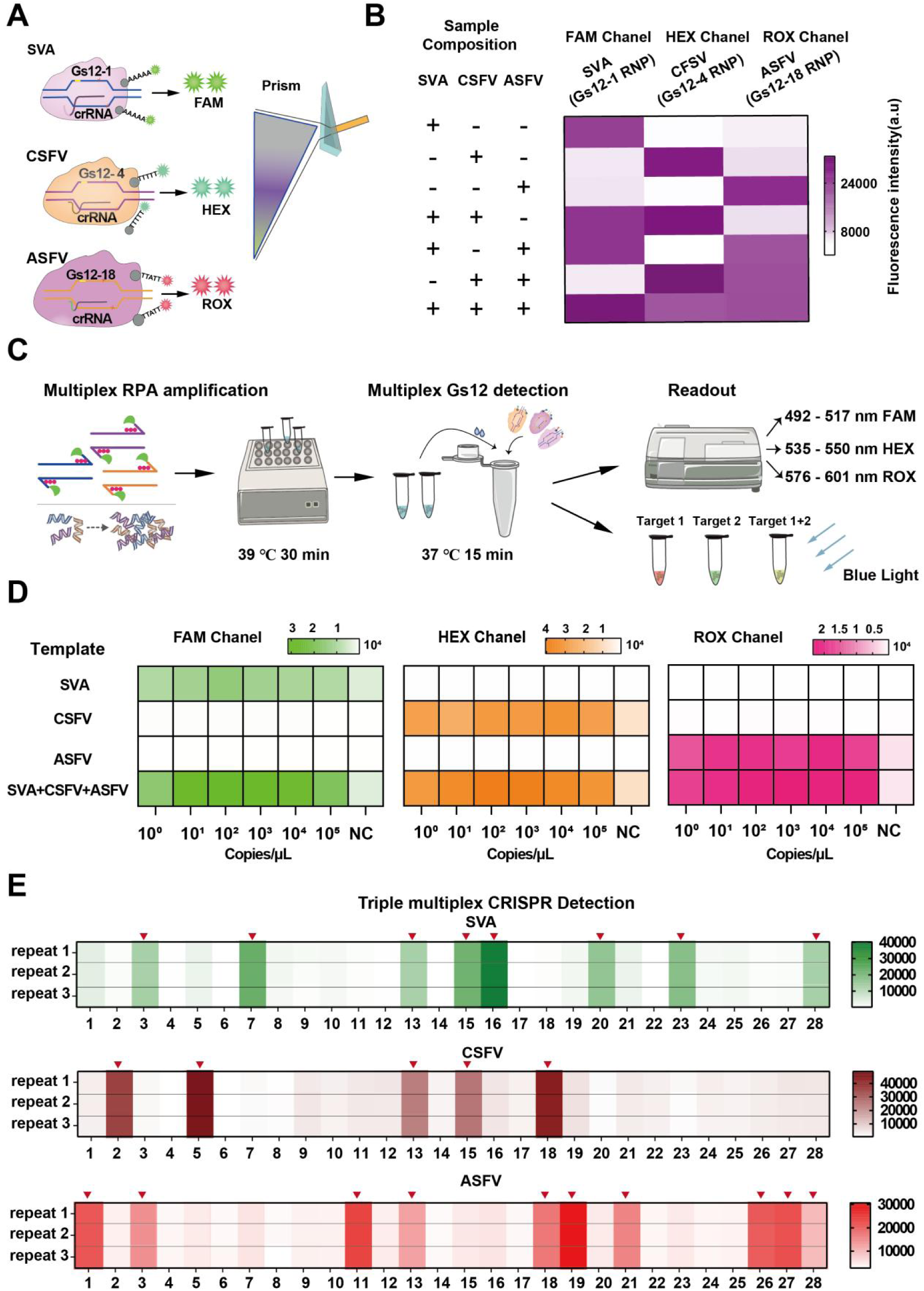
Evaluation of the specificity of detection of three viruses by a multiplexed RPA-CRISPR-Gs12 assay. **A.** Schematic illustration of triple-channel multiplex CRISPR-based nucleic acid detection with Gs12-1, Gs12-4, and Gs12-18 nucleases. ASFV, stands for African Swine Fever Virus. SVA, stands for Seneca Valley Virus. CSFV, stands for Classical Swine Fever Virus. FAM, stands for 6-Carboxyfluorescein, HEX, stands for Hexachloro-fluorescein, ROX, stands for carboxy-X-rhodamine. **B.** In-sample multiplexed detection of ASFV, CSFV and SVA dsDNA with Gs12-1, Gs12-4, and Gs12-18 nucleases. RNP (ribonucleoprotein) refers to a complex consisting of the Gs12 protein and crRNA. The notation +/- indicates whether detection of the target is present (+) or absent (-). **C.** Schematic of CRISPR-Gs12-based nucleic acid detection system coupled with multiplexed RPA for triple or dual target nucleic acid assays. RPA stands for Recombinase polymerase amplification. **D.** Evaluation of CRISPR-Gs12-based multiplexed nucleic acid detection sensitivity for ASFV, SVA, and CSFV using a microplate reader. NC stands for negative control. **E.** Simultaneous detection of ASFV, SVA, and CSFV in a single reaction by CRISPR-Gs12-based multiplexed nucleic acid detection using a microplate reader. Inverted triangle indicated a positive signal. N = 3.

Next, we established a detection process integrating multiplex RPA with the CRISPR-Gs12 system (Figure 6C). Using a simplified boiling method, genomic DNA or RNA extraction from samples precedes RPA or RT-RPA amplification of three target fragments. This rapid amplification was completed within approximately 30 minutes. The amplified products are then introduced into the CRISPR-Gs12 system, designed for ASFV p72, SVA VP1, and CSFV E2 amplicons. The readout was designed to utilize a Microplate Reader or operate under blue light (Figure 6C). We have developed a triple multiplex nucleic acid detection system based on a microplate reader and a dual multiplex CRISPR/Gs12 blue light visual detection assay (Figure S16A). We then evaluated the sensitivity of this assay, and the results showed that the detection limit of each target can reach as low as single copies per μL (Figure 6D and Figure S16B). Subsequently, we simulated 28 clinical samples with single, two and triple ASFV, SVA and CSFV infections. The results demonstrate that our new technique can accurately detect single or multiple targets in a single reaction system, consistent with qPCR detection (Figure 6E, Figure S16C and Figure S17). In conclusion, the results suggested that combining multiplex RPA and CRISPR-Gs12 systems would provide a rapid and sensitive multiplex nucleic acid assay.

## Discussion

Cas12a, a CRISPR-associated nuclease, has broad utility in genome engineering, agricultural genomics, and molecular diagnostic applications^5^. Therefore, identifying novel Cas12a systems with high *cis*- and *tran*s-cleavage activities, broad PAM ranges, or specific ssDNA preferences is crucial. In this study, we explored the evolutionary diversity of CRISPR/Cas12a systems through a combination of bioinformatic screening and experimental characterization. We established a comprehensive framework for *de novo* annotation of Cas12a systems from prokaryotic genomes and subsequently selected the most promising candidates for experimental validation.

Through our comprehensive screen of Cas12a systems, we successfully identified 21 active Gs12 nucleases *in vitro*, with three demonstrating activity in mammalian cells. Notably, Gs12-10 exhibits a level of PAM-less gene editing activity comparable to that of Cas12a editors, while our engineered Gs12-7MAX demonstrated superior editing efficiency compared to previously optimized enAsCas12a-HF1. Furthermore, we developed a powerful multiplexed CRISPR-based nucleic acid detection system that harnesses these newly identified orthogonal Cas12a nucleases. We also designed engineered scaffolds of crRNA for these Gs12 nucleases to enhance the specificity of multiplexed nucleic acid detection. Overall, our study presents a conceptually novel approach to discovering the diversity of the Cas12a family, offering significant potential for expanding the applications of CRISPR-Cas12a-based gene editing and diagnostics.

Interestingly, our screen assay identified the first natural PAM-less Cas12a family member. The ever-expanding set of CRISPR-Cas12a technologies exhibits remarkable flexibility in DNA targeting^4^. This flexibility is a significant advantage but comes with an ever-present constraint: the requirement for a PAM flanking each target^7^. While there are plenty of Cas12a variants that utilize alternative PAMs, such as Mb2Cas12a-RVRR^33,44,45^, and other engineered variants like PiCas12a (with KKYV/TTGS PAM) and Fn3Cas12a (with YTV PAM)^19^, the range is still limited. Ongoing efforts are focused on realizing PAM-free Cas nucleases through natural ortholog mining and protein engineering, such as the near-PAMless engineered SpRY Cas9 variant (with NYN/NRN PAM) was developed^46^. In comparison to other Cas12a, we demonstrate that Gs12-10 possesses an unprecedented capacity to recognize a broad spectrum of PAMs. Notably, Gs12-10 can identify 52 distinct PAM types, which represents a significant 1.8-fold expansion in recognition range compared to the relative LbCas12a PAM in our study. This remarkable ability to target diverse PAMs highlights the vast potential of Gs12-10 as a versatile and powerful tool for precision genome editing and molecular diagnostics (Figure 2). However, to comprehensively profile the PAM preferences of this Gs12-10, an unbiased *in vitro* or *in vivo* PAM determination assay needs to be performed.

A structural basis can be used for altered PAM recognition by engineered CRISPR-Cpf1 or Cas12a^7^. Engineered Cpf1 or Cas12a variants have altered PAM specificities^44^. Nevertheless, the structure of Gs12-10 nuclease remains unknown, which hinders understanding the mechanism by which it achieves PAM-less activity. For gene editing tool development purposes, this PAM-less Gs12-10 nuclease brings an expanded target scope, which can generate CRISPR Gs12-10-based base editors or prime editors. PAM limitations are also a challenge for CRISPR-based diagnostics. Still, this Gs12-10 nuclease has the potential to be used for CRISPR-based detection, which could simplify and expand the scope of CRISPR diagnostics assays. Engineered CRISPR-Cas12a variants can increase activities and improve targeting ranges for gene editing^11^. To enhance the performance of Gs12-7 and Gs12-10, we screened various variants and identified optimized versions, Gs12-7MAX and Gs12-10MAX, which exhibited increased gene editing activity. Notably, we found that Gs12-7E157R exhibits a 1.27-fold increase in editing efficiency over enhanced high-fidelity enAsCas12a-HF1, with cleavage activity reaching up to 42% without cell sorting (Figure 4). However, the PAM of Gs12-7 is similar to AsCas12a, with a primary preference for canonical “TTTV” PAM. Therefore, it is necessary to broaden the PAM range of Gs12-7 in the future.

Sequence-specific endonuclease Cas12-based biosensors have rapidly evolved into a powerful tool for detecting nucleic acids^28^, however, achieving CRISPR-based four-channel nucleic acid detection remains a significant challenge. Our screen yielded 21 active enzymes with robust *trans*-cleavage activity and ssDNA preference. Leveraging this property, we have developed a suite of RNA-guided endonucleases, including Gs12-1, Gs12-4, Gs12-9, and Gs12-18, along with their corresponding structurally engineered crRNAs and ssDNA-FQ reporters. Specifically, our method combines the multiplexed RPA reaction with the specific *trans*-cleavage activity of these Gs12 nucleases, allowing for multi-channel multiplexed detection via target-specific ssDNA-FQ reporters that utilize distinct Gs12 nucleases. As a proof-of-concept, we used this multiplexed CRISPR-based method to simultaneously detect three viruses, each with a detection limit of a single copy per microliter. Our study presents a conceptually novel approach to designing multiplexed CRISPR-based diagnostic systems, offering significant potential for expanding the applications of CRISPR-based diagnostics. Unfortunately, the one-pot RPA-CRISPR-Gs12-based method has yet to be fully established. Therefore, there is still a need to improve our CRISPR-based multiplex nucleic acid detection method in the future.

In conclusion, we explored the evolutionary diversity of CRISPR/Cas12a systems through a combination of bioinformatic screening and experimental characterization. This led to identifying novel CRISPR-Gs12 systems with high *cis*- or *trans*-cleavage activities, specifically those with PAM-less gene editing activity or specific ssDNA preferences. By making our computational and experimental protocols publicly available, we anticipate that this discovery framework will promote even deeper mining and understanding of the natural reservoir of Cas12a systems. The discovery of diverse functions in Gs12 not only provides deeper insight into the CRISPR/Cas12a family but also has the potential to facilitate its expanded application and power the exploration of the undiscovered properties of other CRISPR/Cas systems.

## Materials and methods

### Bioinformatic pipeline for novel Cas12a orthologs

We developed a bioinformatic pipeline to mine novel Cas12a orthologs from microbial genomes. Initially, we downloaded 278,629 genome bins from the Global Microbial Gene Catalog (GMGC) database using its API (https://gmgc.embl.de/api/v1.0/genome_bin/)^36^. We then filtered out genome sequences with lengths greater than 20 kb and scanned them for CRISPR arrays using MinCED (version 0.4.2) software^37^. Next, we employed Prodigal (version 2.6.3) software^37^ to predict open reading frames (ORFs) and translate them into protein sequences within the CRISPR array-containing genomes.

A custom Python script was used to extract ten protein sequences flanking each CRISPR array, which were then clustered at 90% identity using CD-HIT software (version 4.8.1) to generate a protein library. We utilized multiple sequence alignments of Cas proteins from previous studies as templates and annotated the protein library for Cas12a orthologs using psi-BLAST software (BLAST 2.13.0+)^39^. A custom Python script was developed to retrieve CRISPR-Cas gene loci from genome bins, filter them based on the structural architecture of CRISPR-Cas2-Cas4-Cas1-Cas12a, and identify Cas12a as candidates for further investigation. The candidate proteins were searched against the NCBI NR^47^ database using BLASTP (BLAST 2.13.0+)^48^ to identify similar sequences. We took the highest identity hit from the search results for each protein as an indicator of homology with known proteins. A number of Cas12a orthologs were screened for amino acid homology from low to high, and their GMGC IDs and species origin are shown in Supplementary Table 2.

### Prediction of functional domains in Cas proteins

The hh-suite (version 3.3.0) software package was used to predict the functional domains of Cas proteins^49^. A search against the UniRef30 database (https://uniclust.mmseqs.com/)^50, 51^ was performed using hhblits software^50^, and its output was used as input for hhsearch ^52^ software against the Pfam database (https://www.user.gwdg.de/~compbiol/data/hhsuite/databases/hhsuite_dbs/)^49,53^ to obtain functional domains.

The functional domains of Cas12a proteins comprise REC1, REC2, PI, RuvC, and NUC. However, only REC1, RuvC, and NUC have available information in Pfam, allowing us to annotate only these three domains. To annotate the remaining two domains, REC2 and PI, we collected known Cas12a protein sequences and extracted the corresponding domain sequences for multiple sequence alignment. We successfully annotated these two domains using HMMER (version 3.1b2) software ^54^.

### Prediction of protein structures and pairwise comparison

The spatial structures of candidate proteins were predicted using AlphaFold2^40^ in monomer mode. Five structure models were generated for each protein, and the best model (ranked_0.pdb) was selected. We chose five reported Cas12a family members, namely AsCas12a, LbCas12a, FnCas12a, MbCas12a, and PiCas12a, as templates for structure comparison. Using TM-align^41^ (version 20220412) software and PyMOL (version 3.0.0) software (http://www.pymol.org/pymol), we performed a structural alignment between the predicted structures and the templates, obtaining TM-scores and RMSD values to reflect structural similarity. The resulting structures were visualized using PyMOL.

### Phylogenetic analyses

The phylogenetic tree was visualised using the iTOL website^55, 64^. The amino acid sequences of Cas proteins were aligned using MAFFT (version 7.490) software with default parameters^56^. The resulting alignment was then utilized to construct a phylogenetic tree using FastTree (version 2.1.11) software, again with default parameters ^57^.

### Plasmids and oligonucleotides

All Gs12-1 to Gs12-21 endonucleases predicted by prokaryotic expression were synthesized and cloned into the *Nco* I and *Hind* III cleavage sites of pET-28a (+) by GeneScript after bacterial codon optimization. The corresponding amino acid sequences are shown in the Supplementary Table 3. Vectors for eukaryotic expression of Gs12-7 or Gs12-10 were constructed into *Age* I and *BamH* I sites of pLenti-Puro (Addgene plasmids #39481)^58^ following human codon optimization. CrRNA eukaryotic expression vectors were ligated by primer annealing into the modified Gs12 scaffold from Addgene (#67975) in the vector pLenti-U6-Gs12-crRNA-zsGreen. And all crRNA sequences used are all shown in Supplementary Table 4. The annealing primers are listed in the Supplementary Table 3. Gs12-7 or Gs12-10 variants were primed to introduce rapid, targeted mutations using the Fast Site-Directed Mutagenesis kit (TIANGEN). The sequences of all primers used in this study are provided in the Supplementary Table 5.

### Protospacer adjacent motif depletion assay

To generate the plasmid library, we synthesized a ssDNA oligonucleotide (GenScript) containing a random PAM sequence of 7 nts at the 5’ end and a 31 bp target fragment at the 3’ end. Following PCR amplification with oligo-F and oligo-R primers (Supplementary Table 5), the plasmid was assembled into the pUC19 plasmid by homologous recombination to form a random PAM library^4^. The plasmid library was transformed into *E. coli* DH5α, and >10^7^ cells were collected, and the plasmid was extracted (OMEGA). We cloned the crRNA sequence and 21 Gs12 nucleases and LbCas12a into pACYC-Duet-1 from MiaoLing Biology (#P0049), respectively.

Subsequently, 100 ng of the above plasmid libraries were transfected into control *E. coli* receptor cells (Chl^+^) containing pACYC-crRNA-Gs12-1-21, pACYC-crRNA-LbCas12a or control pACYC-Duet-1 plasmids (Chl^+^). Transfected *E. coli* was then screened on LB medium containing dual antibiotics (34 µg/mL chloramphenicol and 100 µg/mL ampicillin) and incubated at 37 °C for 16 h. Plasmid DNA was extracted from the screened *E. coli* (OMEGA). Amplicon sequencing libraries were generated by polymerase chain reaction (NEB) (Supplementary Table 3). The corresponding PAM sequences were analyzed by high-throughput sequencing^4^, and the corresponding WebLogs were generated.

### Analysis of Protospacer adjacent motif

To identify the 7-bp random sequence, we analyzed sequences flanking the target sequence. We normalized the frequency of each PAM sequence by dividing its read count by the total read count of all PAM sequences. Following normalization, we calculated the base-2 logarithmic fold change of each PAM between the experimental and negative control groups. We considered a base-2 logarithmic fold change exceeding 3.5 relative to the negative control as significant. The PAMs with significant cleavage were extracted and visualized using WebLogo (https://weblogo.berkeley.edu/logo.cgi)^59^.

### Protein expression and purification of Cas12a

We synthesized Gs12 nucleases with *E. coli* codon optimization (GenScript) on a pET-28a plasmid (Supplementary Table 3). The bacterial expression vectors were transformed into BL21(DE3) chemosynthetic cells (AngYuBio). We grew 10 mL of starter culture in Luria-Bertani (LB) medium at 37°C overnight, then inoculated it into 2L LB medium and cultured it at 37°C and 200 rpm until the OD600 reached 0.6. At this point, we induced protein expression with IPTG (Isopropyl-beta-D-thiogalactoside) (Sigma) at a final concentration of 0.5 mM, cooled the cells to 16°C, and allowed them to express for 16 hours. Subsequently, we harvested the cells by centrifugation at 13,000 rpm for 30 minutes at 4°C and resuspended them in lysis buffer (500 mM NaCl, 20 mM HEPES, 20 mM Tris, 10 mM imidazole, 1 mM DTT, 1% Triton X-100, 0.005 mg/mL lysozyme, pH = 7.5).

We then performed 10 cycles of high-power sonication (ScientZ; China) and clarified the lysate by centrifugation at 10,000 × g for 30 minutes. The supernatant was pooled and loaded onto a gravity column packed with Ni-NTA agarose (Qiagen) and incubated for 1 hour at 4°C. We washed the column twice with rinse water and then with elution buffer (500 mM NaCl, 20 mM Tris, 50 mM imidazole, pH = 7.5), followed by elution with elution buffer (500 mM NaCl, 20 mM Tris, 300 mM imidazole). We analysed the eluted proteins by SDS-PAGE and concentrated them to 500 μL before applying them to an AKTA pure protein purification system (Cytiva) on a Superdex 200 increase 10/300 GL column (Cytiva). The peak fractions were concentrated to 100 μL, and protein concentrations were determined using a DS-11 FX+ spectrophotometer (DeNovix) before storing them at -80°C.

### Preparation of crRNA

To generate crRNAs, we designed upstream primers with a T7 promoter and scaffold sequences. In contrast, downstream primers contained a 15 bp sequence complementary to the upstream primer and a 20/23 bp targeting sequence (Supplementary Table 5). We performed PCR amplification and collected the products using Qiagen’s protocol. Typically, 20 µL of reaction mixture containing 1 µg of template was prepared for *in vitro* transcription using the HiScribe T7 High Yield RNA Synthesis Kit (NEB) and incubated overnight at 37°C. We treated the crRNA with 4 U DNase I (NEB) to remove any remaining DNA templates for 15 minutes. Subsequently, RNA was purified using the Monarch RNA Cleanup Kit (NEB) according to the manufacturer’s instructions. All crRNA sequences used in this study are listed in Supplementary Table 5.

### In vitro *cis*-cleavage assay of Cas12a

We performed an *in vitro cis*-cleavage assay using purified Gs12-1 to Gs12-21 and LbCas12a proteins at 37°C in rNEBuffer 2.1 cleavage buffer for 30 minutes. To assess the activity of the Gs12 protein *in vitro*, we varied the reaction time to 0.5, 2, 10, and 20 minutes, respectively. The cleavage assay consisted of 250 nM Gs12 or LbCas12a proteins, 500 nM of *in vitro* transcription of crRNA, and 200 ng of target DNA. The target DNA was generated by PCR amplicons of p72 (Supplementary Table 8). We added 1 μL Proteinase K (Beyotime) to the cleavage assay and incubated it for 10 minutes at 55°C. Finally, we ran the reactions on a 2% Agarose Gel (BIOWEST). The assessment of cleavage activity was calculated from the grey values of the bands.

### Assessment of gene modification by T7E1 cleavage assay and high-through sequencing

To evaluate the gene modification activity of Gs12 protein in HEK293T cells, we transfected 500 ng of Gs12 and 250 ng of crRNA expression plasmids into a 24-well plate using jetPRIME transfection reagent (Polyplus). Genomic DNA (gDNA) was extracted approximately 72 hours post-transfection using the TIANamp Genomic DNA Kit (TIANGEN). We then amplified genomic loci by PCR using DNA Polymerase (NEB) with approximately 200 ng of gDNA as a template and the primers listed in Supplementary Table 5.

For T7E1 cleavage assay-based gene modification assessment, we denatured, annealed, and digested 200 ng of purified PCR products with T7EI (NEB) at 37°C for 30 minutes following the instructions^60^. We added 1μL Proteinase K (Beyotime) to the cleavage assay and reacted for 10 minutes at 55°C. Finally, we ran the reactions on a 2% Agarose Gel (BIOWEST). The assessment of gene modification activity was calculated from the grey values of the bands. For high-through sequencing-based gene modification assessment, we amplified genomic loci by PCR. We added adapters using DNA Polymerase (NEB) with approximately 200 ng of gDNA as a template. The primers are listed in Supplementary Table 5. We then sequenced the PCR products on an Illumina MiSeq Sequencer. Nucleotide substitutions and Indels were analyzed using CRISPResso2^61^.

### Evaluation of predicted off-targets of Cas12a

To assess potential off-target effects, we used CRISPR-Offinder to predict possible off-target genes for the transfected crRNA sequences^62^. We then designed PCR primers to amplify the corresponding off-target sites (Supplementary Table 6). Genomic DNA was extracted from cells 72 hours after transfection and used as a template for PCR amplification of the target genes near the predicted off-target sites. The resulting amplicons were subjected to high-throughput sequencing. The activity of predicted off-target sites was subsequently analyzed using CRISPResso2^61^.

### The evaluation of *in vitro trans*-cleavage characteristics of Cas12a

The *in vitro trans*-cleavage activity of Gs12 was assessed by detecting fluorescence signals in a 20 μL reaction system. This system consisted of 250 nM Gs12s, 500 nM corresponding crRNA, 400 nM ssDNA-FQ reporters (Synthesised by Tyhygene) (Supplementary Table 7), 500 nM dsDNA target, and 1×CutSmart Buffer (ShangruiBio, WuHan). The dsDNA targets were PCR-amplified 750 bp fragments from a previously constructed ASFV p72-pMD18T plasmid^28^. Negative controls were prepared without the dsDNA target. All reactions were incubated at 37°C for 15 minutes, followed by inactivation at 98°C for 2 minutes. Fluorescence detection was performed using an EnSpire microplate fluorescence reader with excitation/emission wavelengths of 576 or 601 nm.

### Fluorescent cleavage assay

The assay was performed in a 20 µL system consisting of 250 nM Gs12s (Gs12-1, Gs12-4, and Gs12-18), 500 nM crRNAs, 500 nM dsDNA targets, and 400 nM ssDNA-FQ reporters (ROX-modified, HEX-modified, or FAM-modified) (Supplementary Table 7). The reaction was incubated at 37°C for 15 minutes. Fluorescence detection was performed using an EnSpire microplate fluorescence reader with excitation/emission wavelengths of 576/601 nm for ROX, 535/550 nm for HEX, and 492/601 nm for FAM. All crRNAs used in this study are listed in Supplementary Table 4.

To evaluate the temperature range for Gs12s proteins, the assay was performed using a 20 µL system consisting of 250 nM Gs12s (Gs12-1, Gs12-4, and Gs12-18), 500 nM crRNAs, 500 nM p72 dsDNA target, and 400 nM ROX-modified ssDNA-FQ reporter. The reaction was incubated at temperatures ranging from 16°C to 65°C for 15 minutes each. Fluorescence detection was performed using an EnSpire microplate fluorescence reader with excitation/emission wavelengths of 576/601 nm. To screen the scaffolds corresponding to different Gs12s proteins, the reaction system consisted of a 20 µL system containing 250 nM Gs12s (Gs12-1, Gs12-4, and Gs12-18), 500 nM scaffold-modified crRNAs (Supplementary Table 4), 500 nM p72 dsDNA target, and 400 nM ROX-modified ssDNA-FQ reporter (Supplementary Table 7). The reaction was incubated at 37°C for 15 minutes. Fluorescence detection was performed using an EnSpire microplate fluorescence reader with excitation/emission wavelengths of 576/601 nm.

### Nucleic acid preparation

A 527 bp fragment of the SVA VP1 gene (GenBank: KT007137.1) and a 507 bp fragment of the CSFV E2 gene (GenBank: MW368990.1) were synthesised by Tsingke and cloned into the pMD18T plasmid. The ASFV p72 fragment was prepared as previously described (GenBank: MK333180.1) (Supplementary Table 8). Two virus co-infection samples and three virus co-infection samples were generated by mixing clinical samples infected with each ASFV, SVA, and CSFV viruses alone to prepare co-infection samples. The QuickExtract DNA Extraction Kit (Lucigen) extracted nucleic acids from all infected samples. An appropriate amount of tissue or blood was added to 80 µL of reagent solution for extraction and reacted at 65 °C for 5 minutes, followed by a 5 minute incubation at 95 °C.

### Detection of targets by qPCR or qRT-PCR

The method for detecting African swine fever virus (ASFV) by quantitative polymerase chain reaction (qPCR) was as previously reported^63^. For the detection of classical swine fever virus (CSFV) or Seneca virus A (SVA), qPCR or RT-qPCR was performed using NovoStart SYBR High-Sensitivity qPCR SuperMix (Novoprotein). The cycling conditions for the CSFV assay consisted of an initial 60-second denaturation step at 95°C, followed by 35 cycles of 20 seconds at 95°C and 45 seconds at 60°C, and finally a 5-minute incubation at 40°C. Fluorescence data were collected during the 65°C annealing step of each cycle.

For the SVA assay, an additional Luna WarmStart RT Enzyme Mix (NEB) was added at the beginning of the procedure to perform the reverse transcription process. The reaction mixture was incubated at 55°C for 20 minutes, followed by the cycling conditions described above. All assays were performed using the CFX real-time PCR assay (Bio-Rad). The primer sets used are listed in Supplementary Table 5. The sensitivity and linearity of the qPCR or RT-qPCR assay were estimated by constructing standard curves. Results with quantification cycle (Cq) values greater than 30 were considered negative. The limit of detection (LOD) was calculated by determining the lowest concentration, which gave a positive result.

### Triple multiplex detection of CRISPR-Gs12 coupled to RPA

The platform combines multiple RPA and CRISPR/Gs12s detection methods. The RPA pellet was resuspended in 29.5 μL of rehydration buffer from the TwistAmp basic RPA kit (TwistDx). The multiple RPA or RT-RPA reactions consisted of 0.4 μL RPA-CSFV-E2-F/R (10 μM), 0.3 μL RPA-ASFV-p72-F/R (10 μM), 0.5 μL RPA-SVA-VP1-F/R (10 μM) (Supplementary Table 5), 14.75 μL resuspended RPA solution, 0.6 μL RNA reverse transcriptase (100 U/mL), 3 μL nuclease-free water, 3 μL sample template, and 1.25 μL MgAc (280 mM), with a total volume of 25 μL. The reaction was conducted at 39°C for 30 minutes in a NaCha MultiTemp Platform (Monad).

The CRISPR/Gs12s reaction mixture consisted of 250 nM each of Gs12-1, Gs12-4, and Gs12-18, 500 nM each of their respective crRNAs, 400 nM each of ROX-modified ssDNA-FQ reporter, HEX-modified ssDNA-FQ reporter, and 250 nM of FAM-modified ssDNA-FQ reporter (Supplementary Table 7), 2 μL CutSmart buffer (10×), 2 μL multiplexed RPA amplification product, and a total reaction volume of 20 μL at 37°C for 15 minutes and 98°C for 2 minutes. The primer sets are shown in Supplementary Table 5, and all ssDNA-FQ reporters are listed in Supplementary Table 7.

### RPA reaction mixture preparation and dual multiplex CRISPR-Gs12 visual detection

The RPA pellet was resuspended in 29.5 μL of rehydration buffer from the TwistAmp basic RPA kit (TwistDx). The multiple RPA or RT-RPA reactions consisted of 0.3 μL RPA-ASFV-p72-F/R (10 μM), 0.5 μL RPA-SVA-VP1-F/R (10 μM), 14.75 μL resuspended RPA solution, 0.6 μL RNA reverse transcriptase (100 U/mL), 4 μL nuclease-free water, 3 μL sample template, and 1.25 μL MgAc (280 mM), with a total volume of 25 μL. The reaction was conducted at 39°C for 30 minutes in a NaCha MultiTemp Platform (Monad).

The Dual Gs12s reaction mixture consisted of 250 nM each of Gs12-1 and Gs12-18, 500 nM each of their respective crRNAs, 400 nM each of ROX-modified ssDNA-FQ reporter and 250 nM of FAM-modified ssDNA-FQ reporter, 2 μL CutSmart buffer (10×), 2 μL multiplexed RPA amplification product, and a total reaction volume of 20 μL at 37°C for 15 minutes and 98°C for 2 minutes. The primer sets are shown in Supplementary Table 5, and all ssDNA-FQ reporters are listed in Supplementary Table 7. After the reaction, PCR tubes containing 20 μL of CRISPR/Gs12s reaction were placed on a blue light transilluminator (ShangruiBio, WuHan). The results were recorded using a smartphone.

### Statistical Analysis

Statistical analysis was performed using the R programming language. Mean values ± standard error of the mean (SEM) were calculated for each treatment group in separate experiments. A two-tailed Student’s t-test was employed to determine significant differences between treatment and control groups. P values are reported according to GraphPad style, where: not significant (ns), *P* > 0.05; \**P* < 0.05; \*\**P* < 0.01; *** *P* < 0.001; \*\*\*\**P* < 0.0001.

## Acknowledgements

This work was supported by the Foundation for Innovative Research Groups of the National Natural Science Foundation of China (Grant No.32221005), the National Key Research and Development Program of China (Grant No.2023YFF1000900), and the Major Project of Hubei Hongshan Laboratory (Grant No.2021hszd003 and 2022hszd023), and the STI2030-Major Projects (Grant No.2023ZD0404301-01). We thank Jing Xu and Yeting Wu from the Public Instrument Center of the College of Animal Science & Technology and Veterinary Medicine. We also thank the research group of GMGC for supporting the metagenomic datasets.

## Supplementary Figures

**Figure S1.**
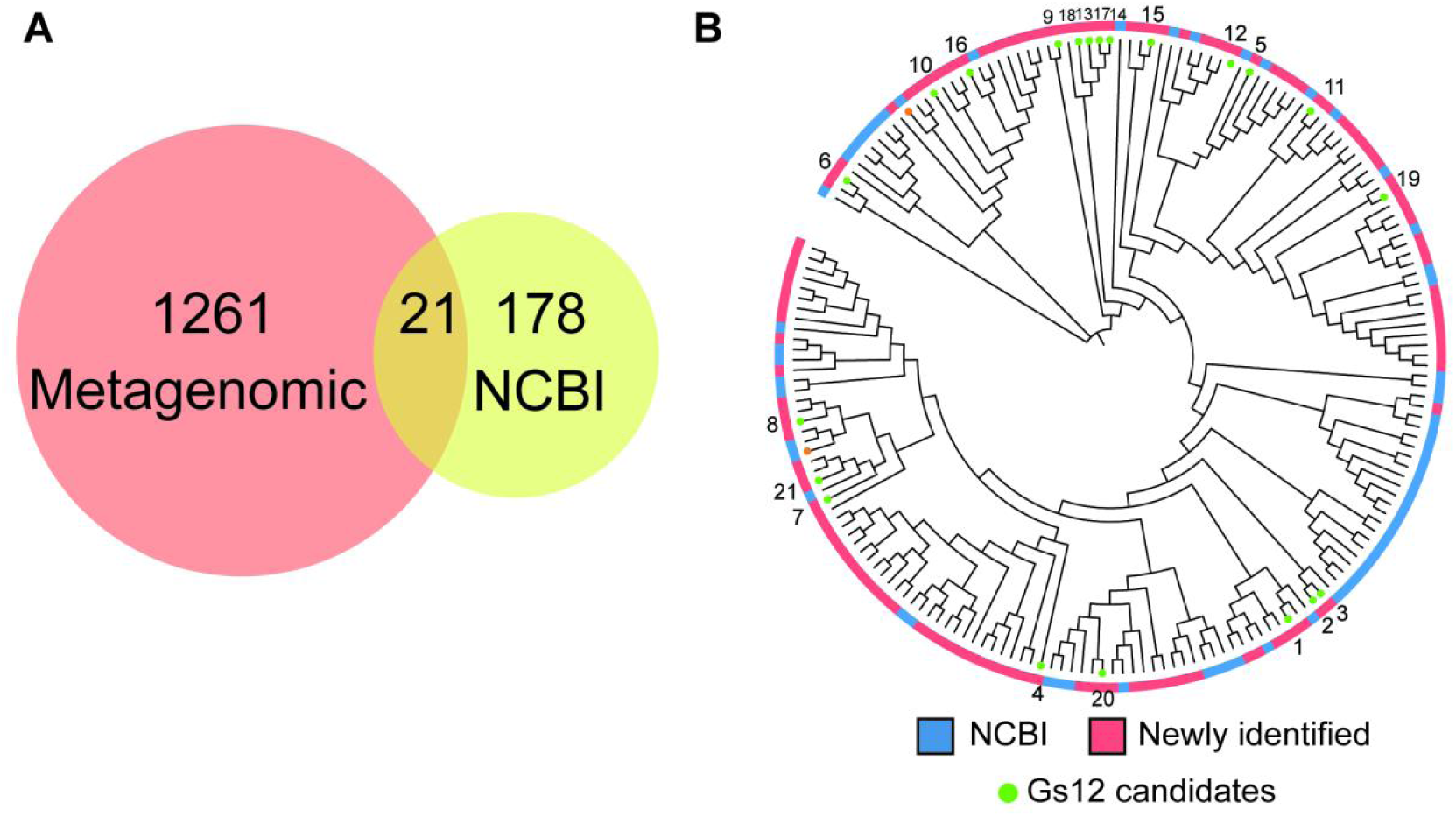
Comparison of the predicted novel Cas12a family members with known Cas12a proteins in NCBI. A. Venn diagram illustrating the relationship between 1,261 newly identified Cas12a loci not previously utilized for genome editing, 178 Cas12a described in National Center for Biotechnology Information (NCBI) database, and the overlapping 21 novel Cas12a endonucleases (Gs12 candidates). B. Phylogenetic tree analysis of the novel Gs12 candidates (green) and their potential classification in the Cas12a subtype family.

**Figure S2.**
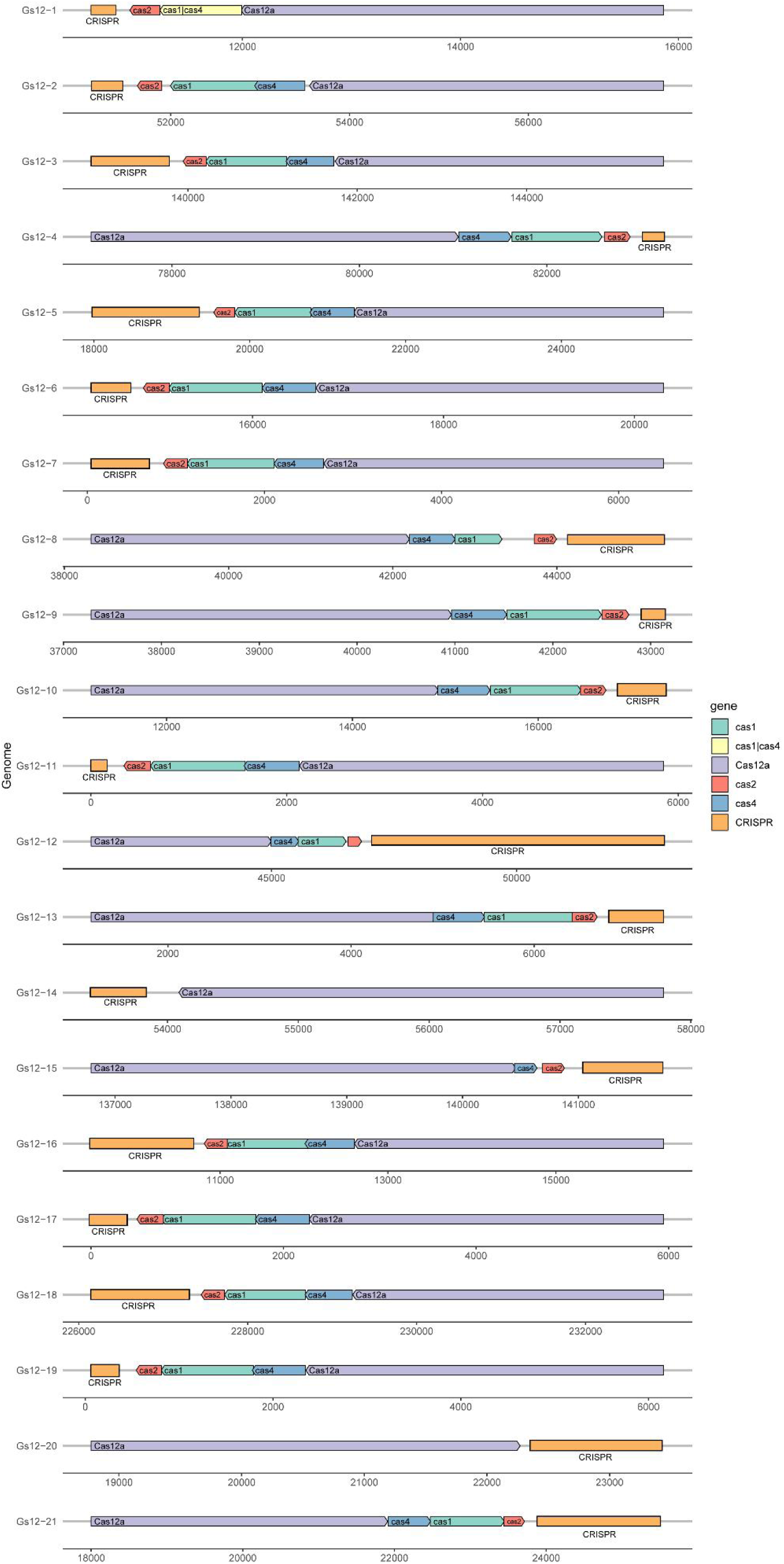
Schematic of the 21 CRISPR-Gs12 array.

**Figure S3.**
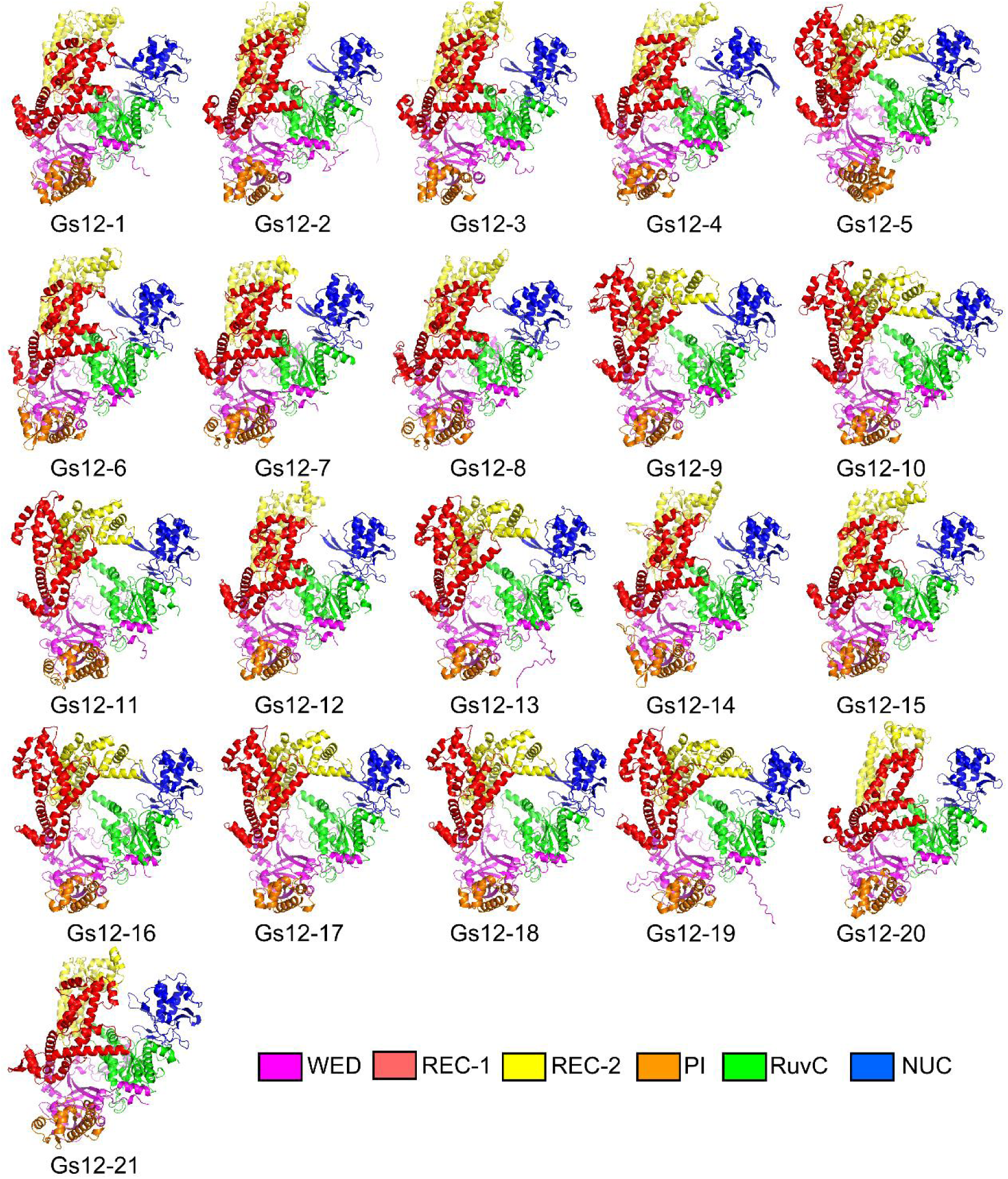
The predicted 3D structure of the 21 Gs12 protein was generated using AlphaFold2. WED stands for wedge domain; REC-1 stands for recognition lobes-1 domain; REC-2 stands for recognition lobes-1 domain; PI stands for PAM-interacting domain; RuvC stands for RuvC domain; NUC stands for nuclease domain.

**Figure S4.**
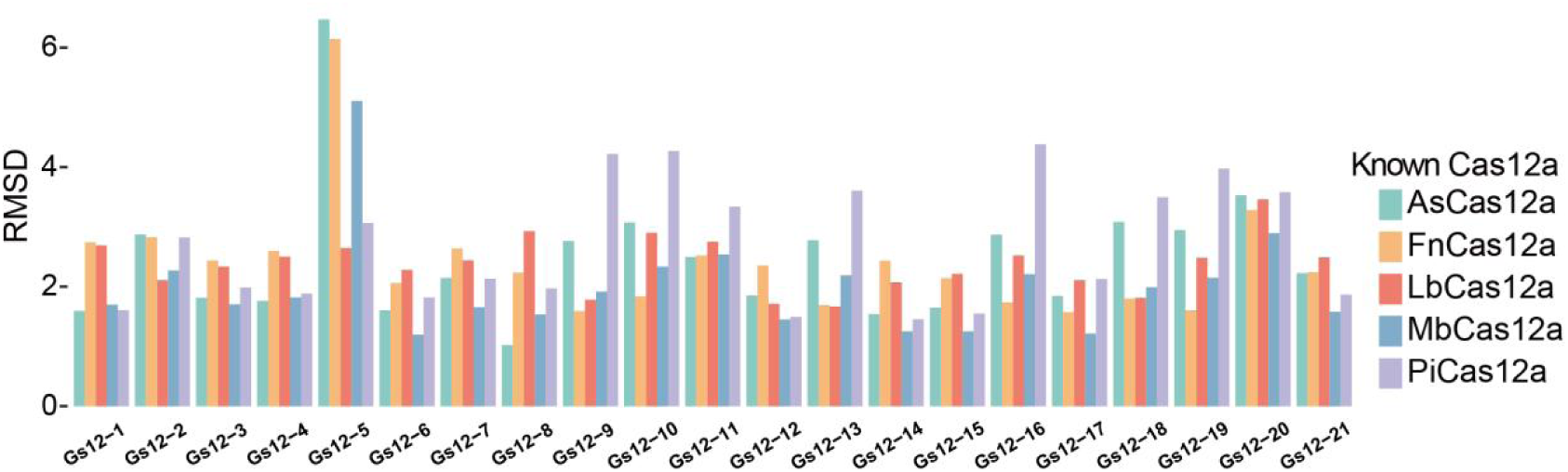
Root Mean Square Deviation (RMSD) values for spatial structure comparisons between 21 Gs12s and classical Cas12a. Note: A lower RMSD value indicates a higher structural similarity between two proteins.

**Figure S5.**
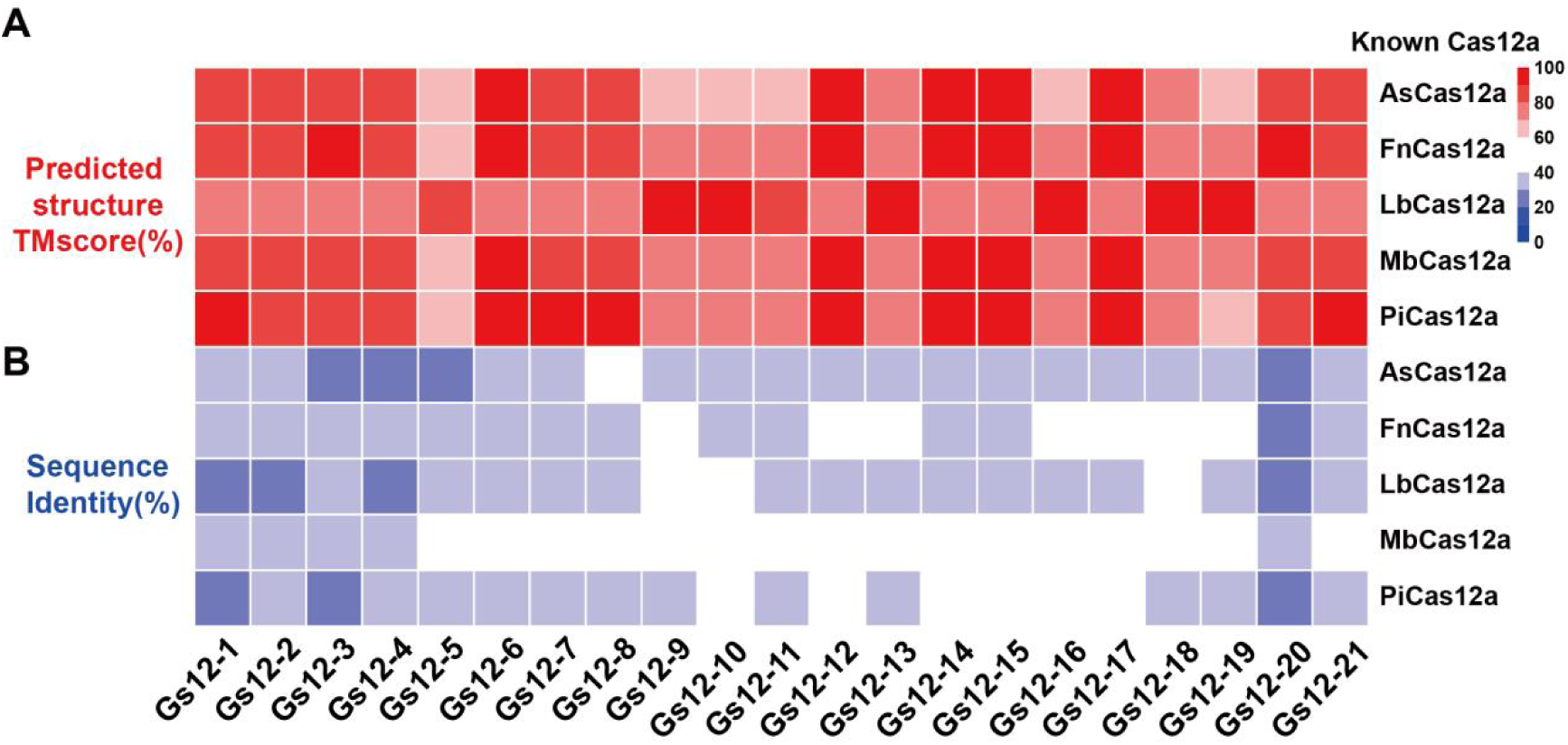
Predicted structure TM scores and sequence identities (%) of 21 Gs12 proteins in comparison to known Cas12a proteins. A. Structure comparisons between Gs12 nucleases and known Cas12a protein pairs using TMalign. B. Amino acid identity (%) of 21 Gs12 nucleases compared to known Cas12a proteins.

**Figure S6.**
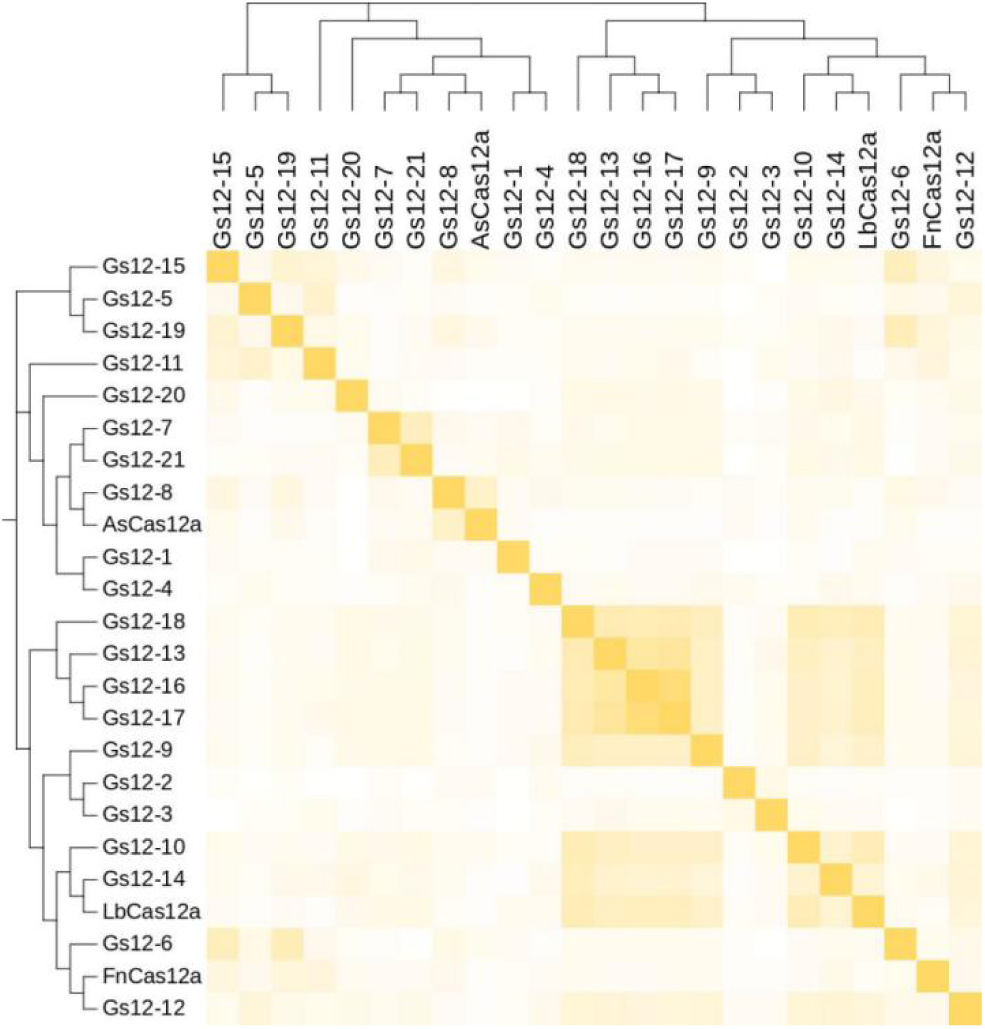
Heatmap of similarity of RuvC structural domains of 21 Gs12s and known Cas12a proteins.

**Figure S7.**
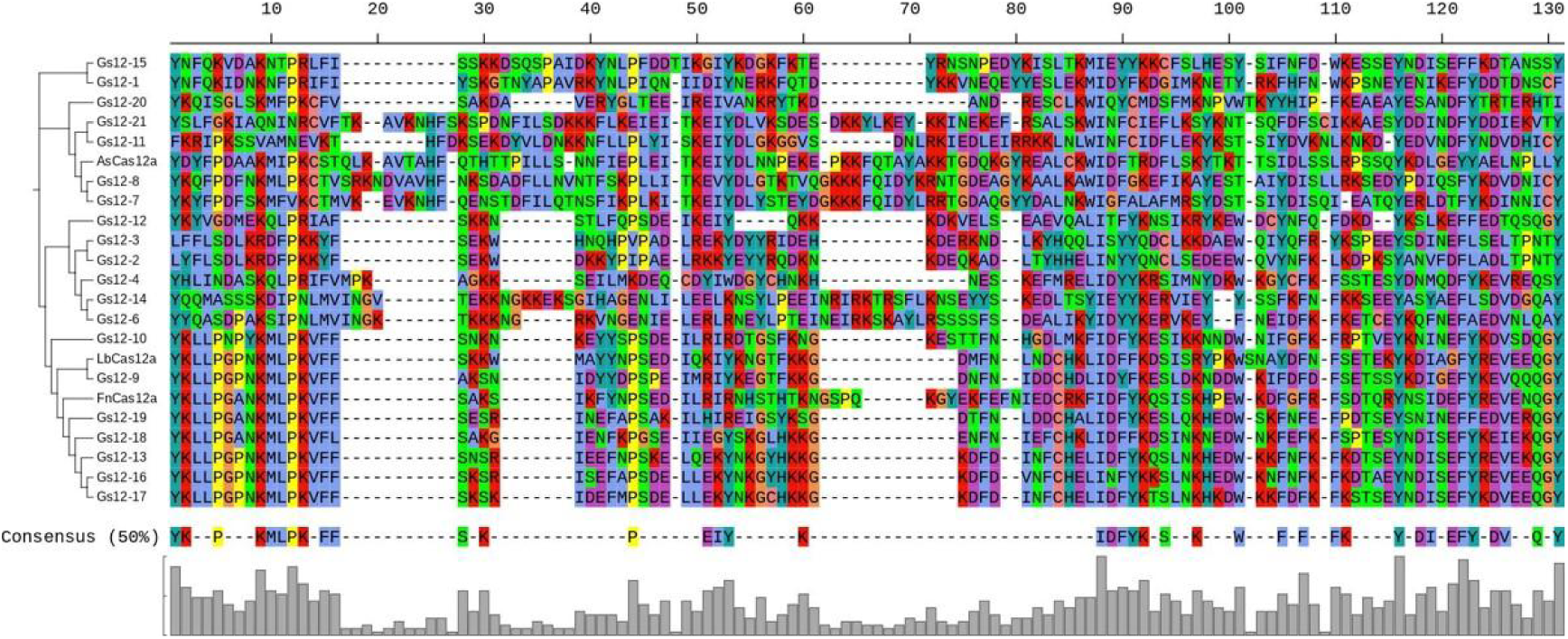
Conservation analysis of PI structural domain of 21 Gs12s.

**Figure S8.**
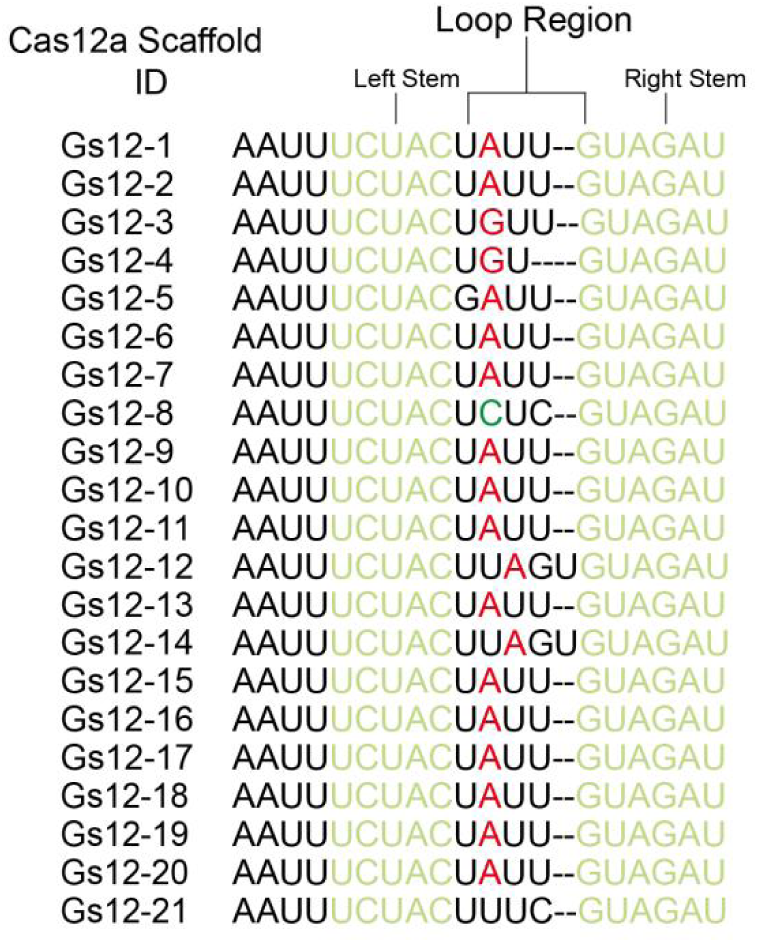
Schematic representation of the scaffold of 21 Gs12 crRNAs.

**Figure S9.**
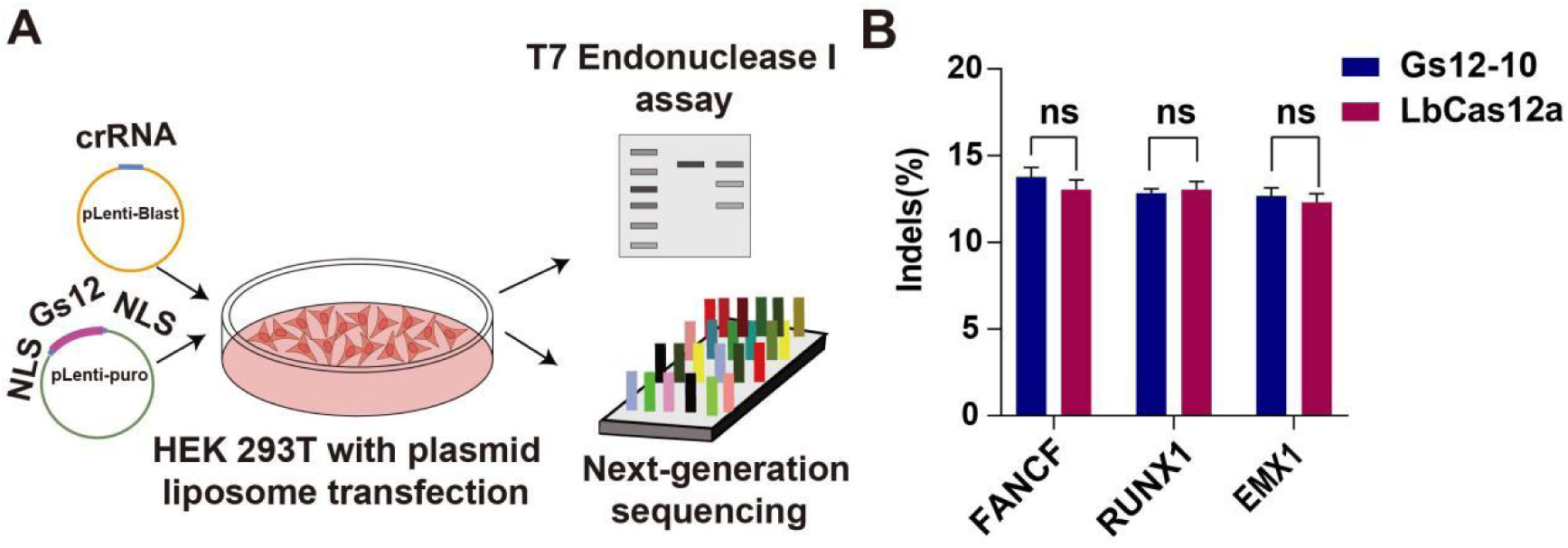
Comparison of the editing efficiency of Gs12-10 and LbCas12a at three endogenous sites. A. Schematic showing the extended targeting range of the Cas12a ortholog in HEK 293T cells transfected with plasmids containing Gs12-10. NLS stands for nuclear localization signals. B. Gene editing efficiency of Gs12-10 and LbCas12a in targeting *FANCF*, *RUNX1*, and *EMX1* genes in HEK 293T. ns: non-significant.

**Figure S10.**
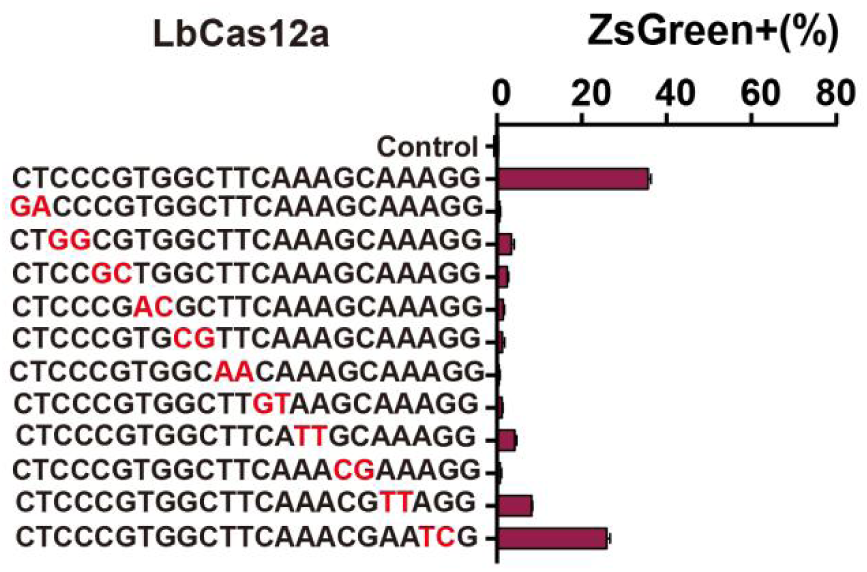
Evaluation of the cleavage ability of LbCas12a for recognize two mismatched target sites using the ZsGreen activation assay. The red letters indicate the presence of two mismatched bases.

**Figure S11.**
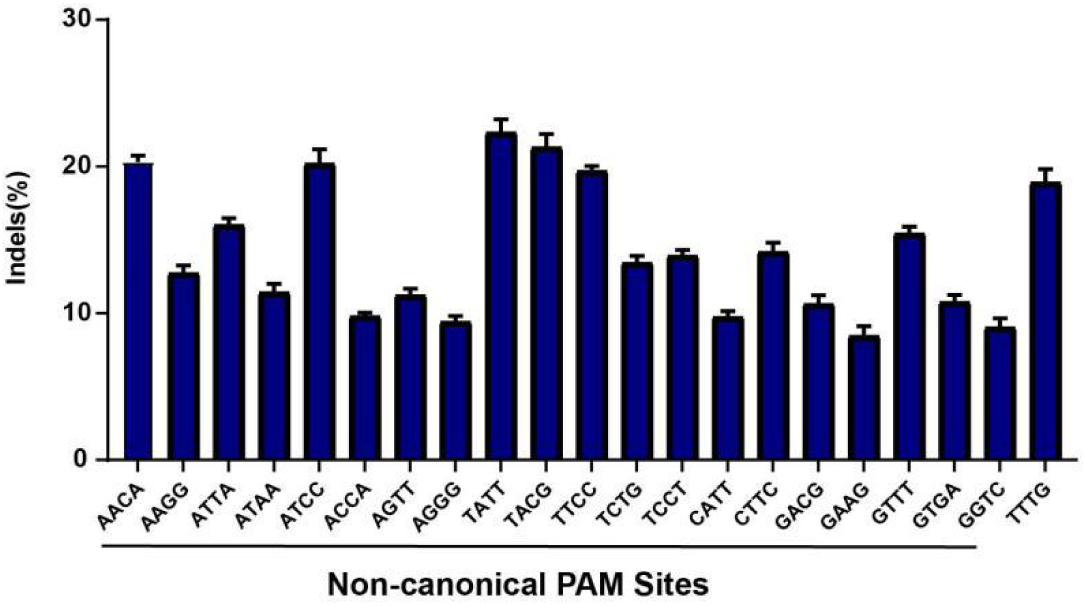
Evaluating the ability of PAM-less Gs12-10 nuclease to cleave DNA at endogenous sites with different PAMs.

**Figure S12.**
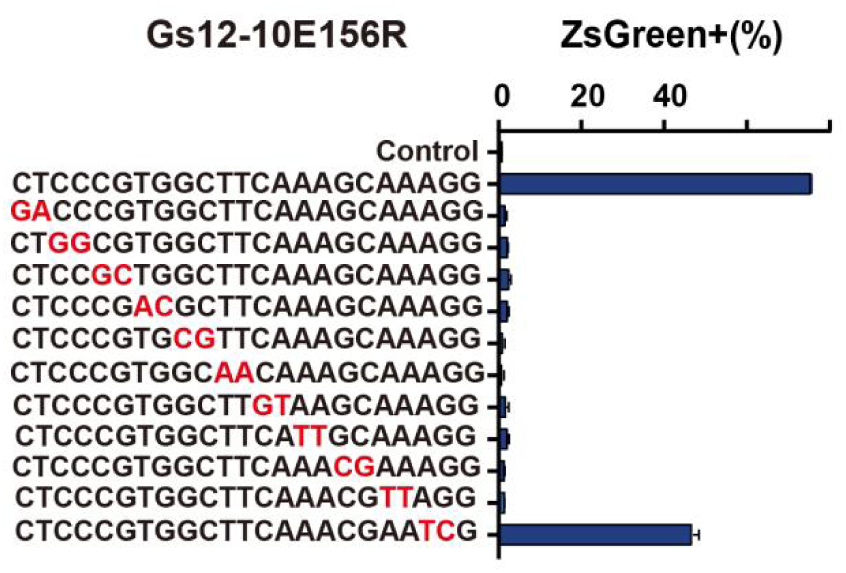
Evaluation of the cleavage ability of Gs12-10E156R for recognize two mismatched target sites using the ZsGreen activation assay. The red letters indicate the presence of two mismatched bases.

**Figure S13.**
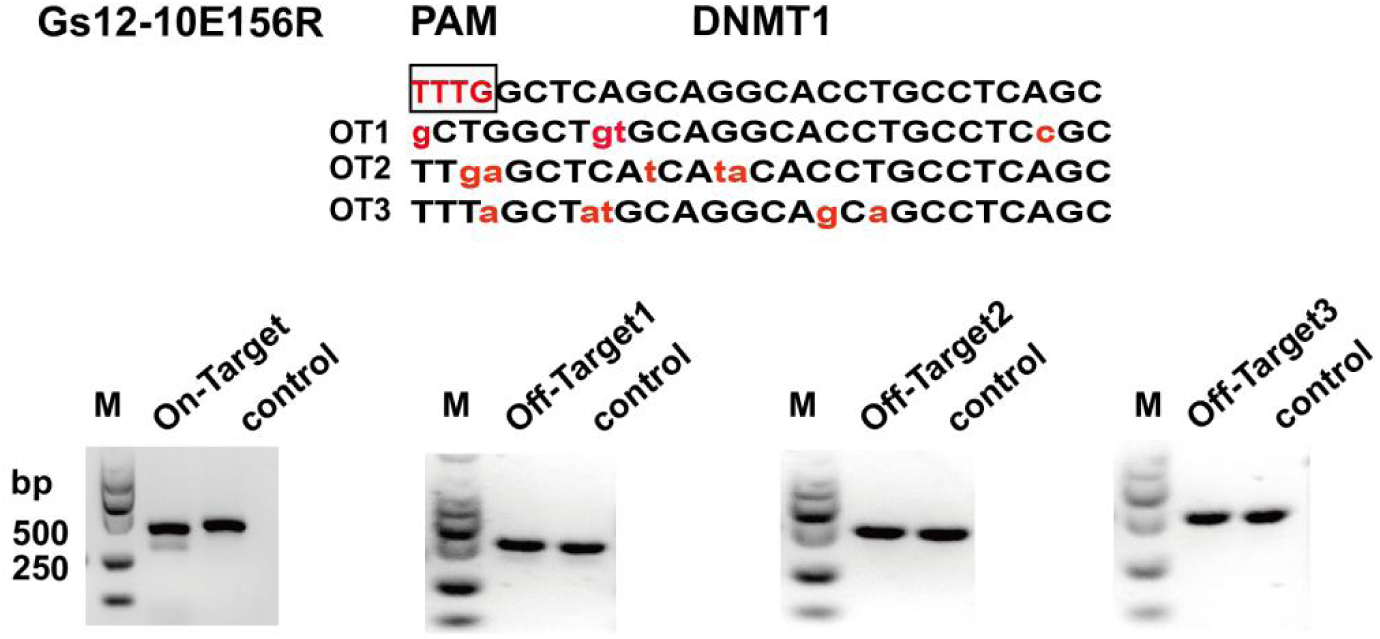
Detection of predicted off-target endogenous sites of Gs12-10E156R (Gs12-10MAX) containing the “TTTG” PAM by using T7E1 cleavage assay.

**Figure S14.**
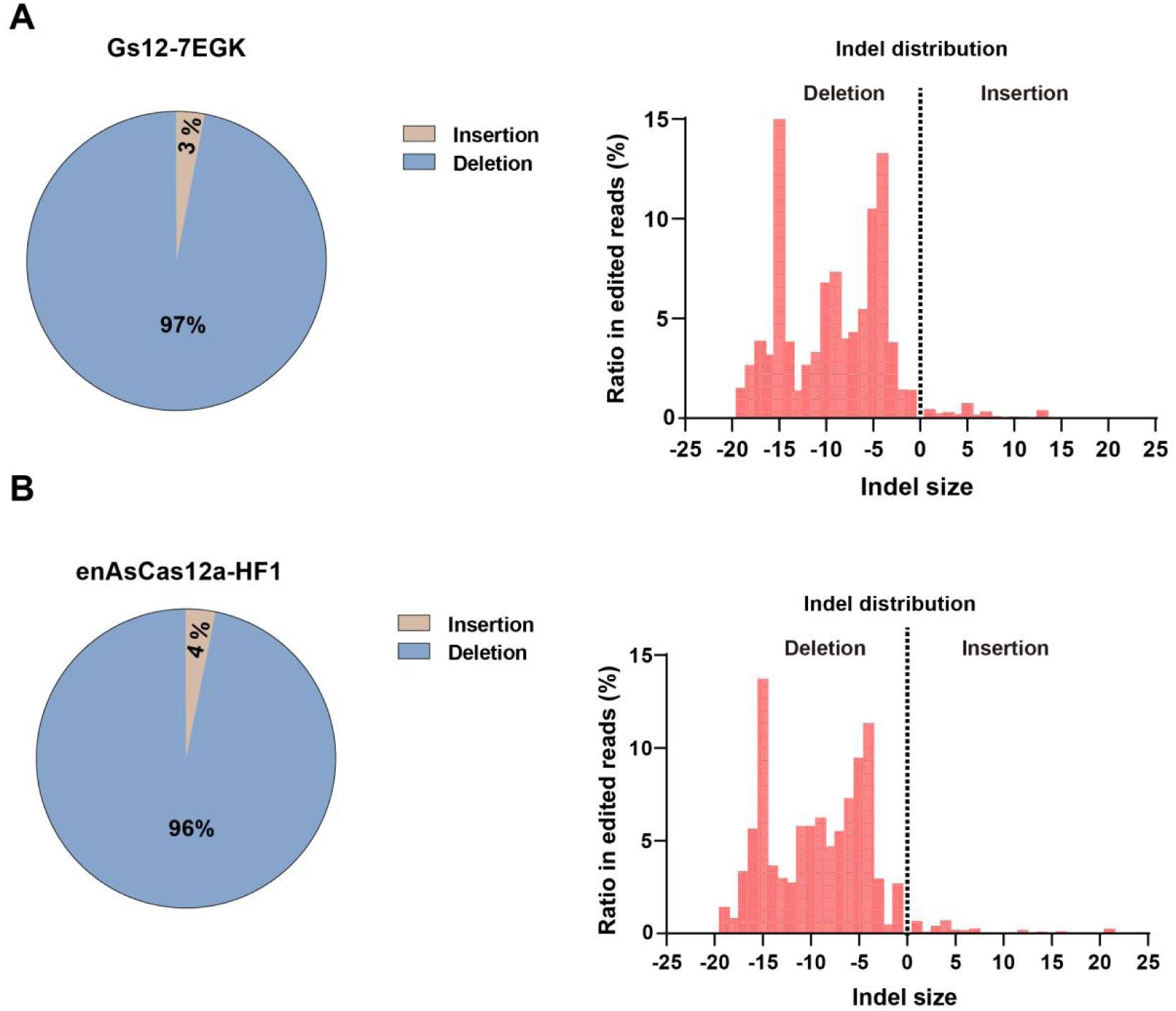
Detecting the outcome of Gs12-7EGK and enAsCas12a-HF1 by high-throughput sequencing. A. The types of gene editing (Left) and Indels distributions (Right) of Gs12-7EGK. B. The types of gene editing (Left) and Indels distributions (Right) of enAsCas12a-HF1.

**Figure S15.**
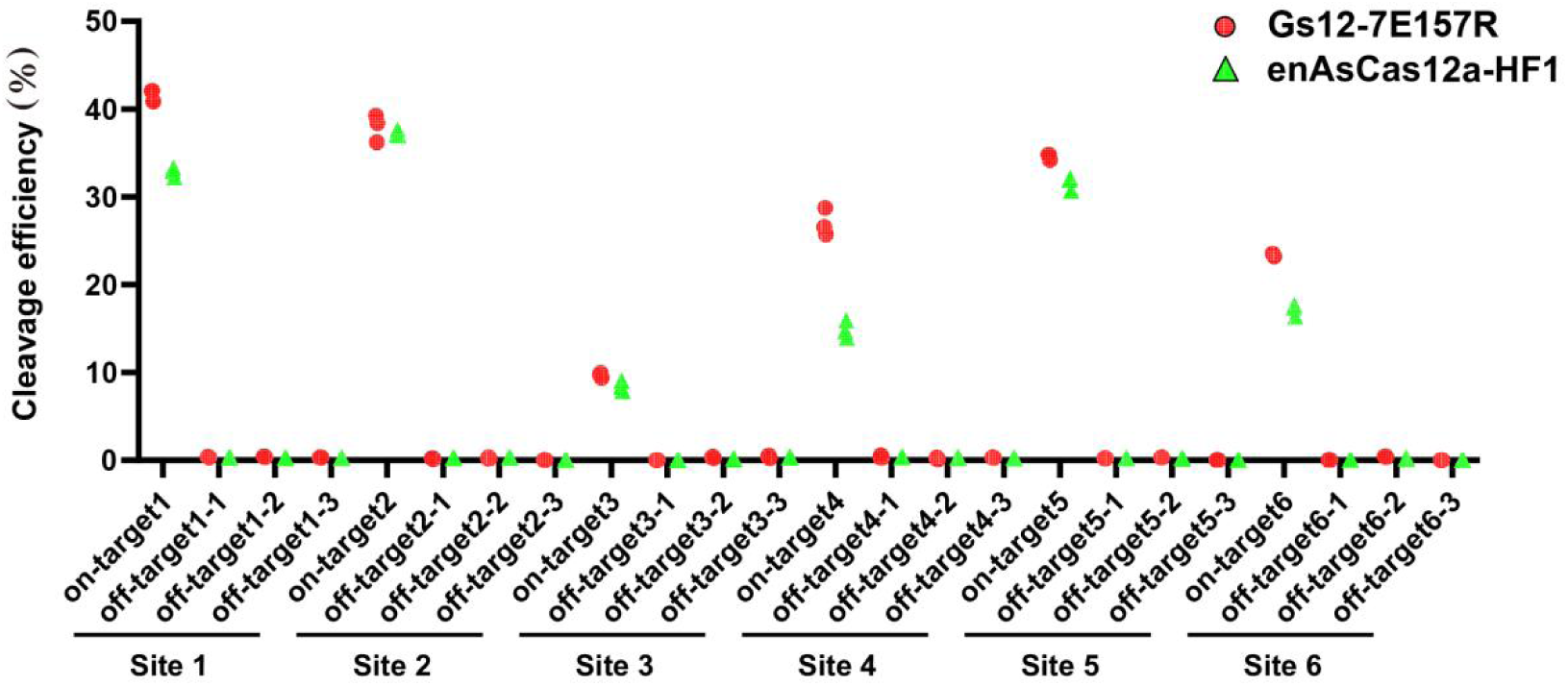
Comparison of the specificity of Gs12-7E157R and enAsCas12a-HF1 for detecting 18 predicted off-target sites corresponding to 6 on-target sites by high-throughput sequencing. Note: Three predicted off-target sites were chosen for each on-target site.

**Figure S16.**
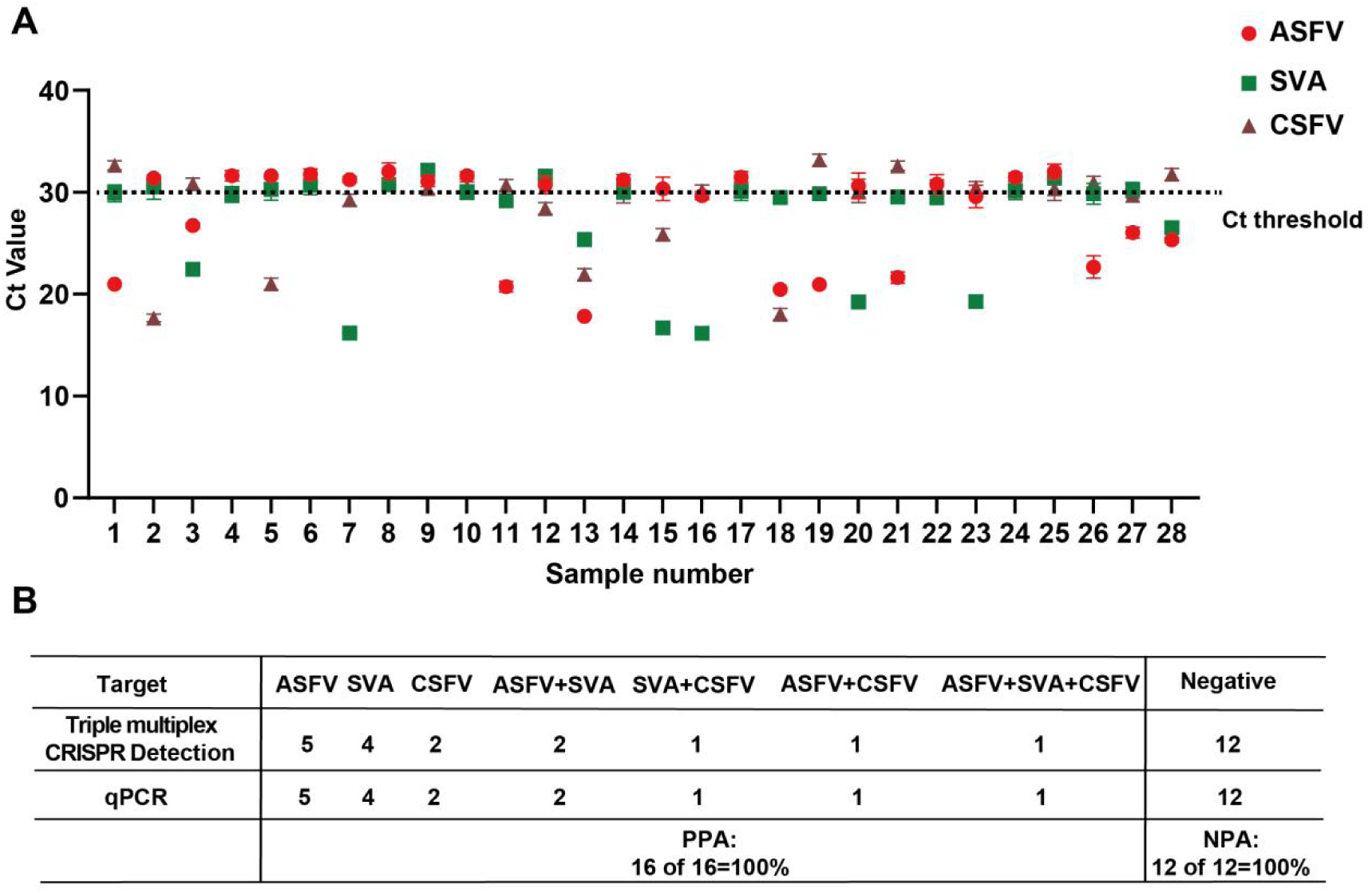
Assessment of Triple multiplex CRISPR/Gs12 nucleic acid assay accuracy. A. Results for 28 simulated clinical samples by qPCR. ASFV, African Swine Fever Virus; SVA, Seneca Valley Virus; CSFV, Classical Swine Fever Virus, ct stands for threshold cycles. B. Statistical analysis of the accuracy of qPCR and multiplex CRISPR/Gs12 nucleic acid assay for the above clinical samples. Mean is shown for n = 3. PPA stands for positive percent agreement; NPA stands for negative percent agreement.

**Figure S17.**
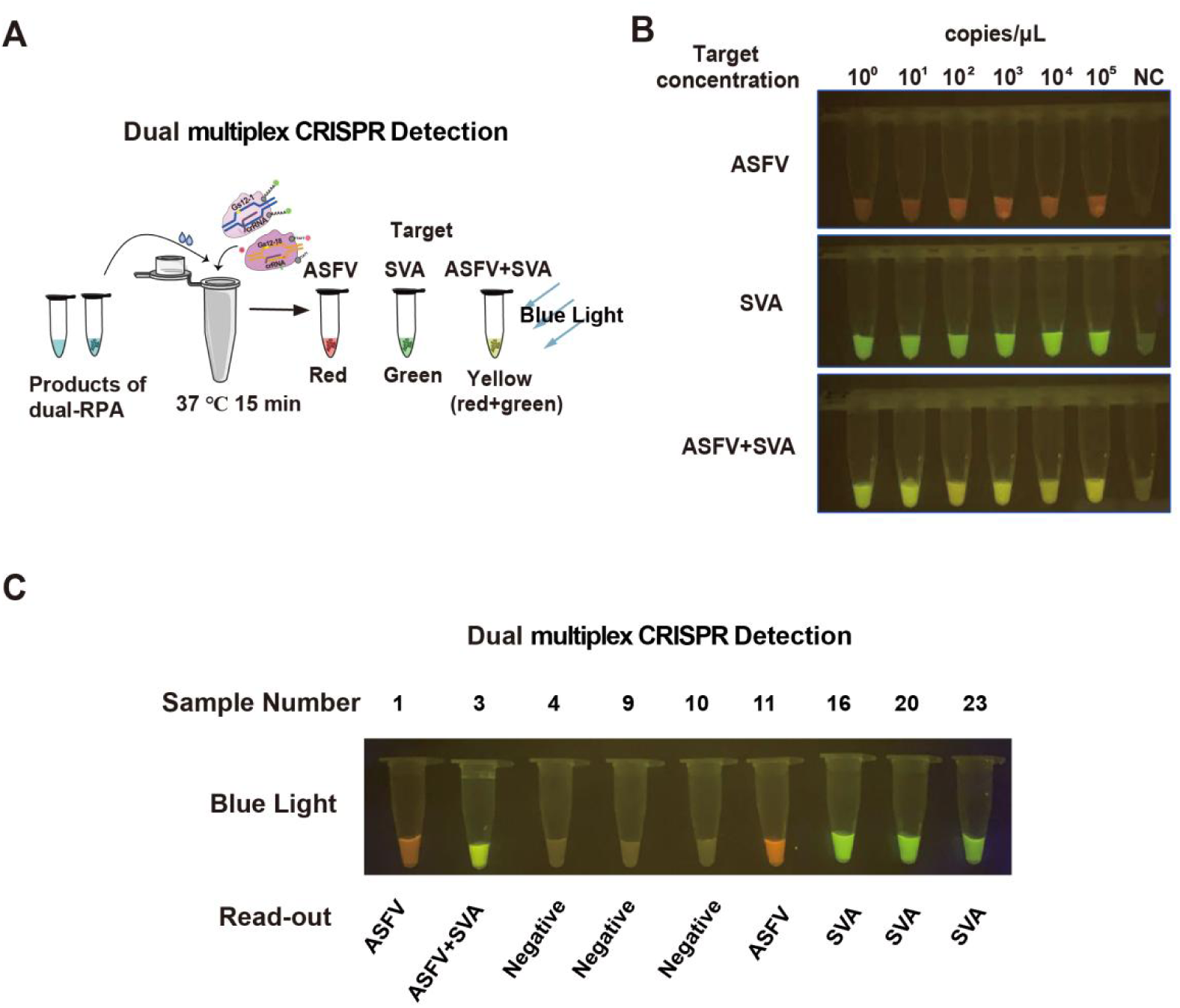
Establishment of a Dual multiplex CRISPR/Gs12 nucleic acid visual detection assay for ASFV and SVA. A. Schematic diagram of the Dual multiplex CRISPR/Gs12 nucleic acid visual detection assay. RPA, Recombinase Polymerase Amplification; ASFV, African Swine Fever Virus; SVA, Seneca Valley Virus; CSFV, Classical Swine Fever Virus. The colors represent the fluorescence signals: Red (degraded ROX-dye ssDNA-FQ reporter) and Green (FAM-dye ssDNA-FQ reporter). B. Assessing the sensitivity of this assay. C. Accuracy testing of the clinical samples listed in Figure S16A.

